# Bacterial swarming reduces *Proteus mirabilis* and *Vibrio parahaemolyticus* cell stiffness and increases β-lactam susceptibility

**DOI:** 10.1101/275941

**Authors:** George K. Auer, Piercen M. Oliver, Manohary Rajendram, Ti-Yu Lin, Qing Yao, Grant J. Jensen, Douglas B. Weibel

## Abstract

Swarmer cells of the gram-negative uropathogenic bacteria *Proteus mirabilis* and *Vibrio parahaemolyticus* become long (>10-100 μm) and multinucleate during their growth and motility on polymer surfaces. We demonstrate increasing cell length is accompanied by a large increase in flexibility. Using a microfluidic assay to measure single-cell mechanics, we identified large differences in swarmer cell stiffness of (bending rigidity of *P. mirabilis*, 5.5 × 10^−22^ N m^2^; *V. parahaemolyticus*, 1.0 × 10^−22^ N m^2^) compared to vegetative cells (1.4 × 10^−20^ N m^2^ and 2.2 × 10^−22^ N m^2^, respectively). The reduction in bending rigidity (∼2-26 fold) was accompanied by a decrease in the average polysaccharide strand length of the peptidoglycan layer of the cell wall from 28-30 to 19-22 disaccharides. Atomic force microscopy revealed a reduction in *P. mirabilis* peptidoglycan thickness from 1.5 nm (vegetative) to 1.0 nm (swarmer) and electron cryotomography indicated changes in swarmer cell wall morphology. *P. mirabilis* and *V. parahaemolyticus* swarmer cells became increasingly sensitive to osmotic pressure and susceptible to cell wall-modifying antibiotics (compared to vegetative cells)—they were ∼30% more likely to die after 3 h of treatment with minimum inhibitory concentrations of the β-lactams cephalexin and penicillin G. The adaptive cost of swarming is offset by the increase in cell susceptibility to physical and chemical changes in their environment, thereby suggesting the development of new chemotherapies for bacteria that leverage swarming for the colonization of hosts and survival.

**Importance:** *Proteus mirabilis* and *Vibrio parahaemolyticus* are bacteria that infect humans. To adapt to environmental changes, these bacteria alter their cell morphology and move collectively to access new sources of nutrients in a process referred to as ‘swarming’. We found that a change in the composition and thickness of the peptidoglycan layer of the cell wall makes swarmer cells of *P. mirabilis* and *V. parahaemolyticus* more flexible (i.e., reduced cell stiffness) and they become more sensitive to osmotic pressure and cell-wall targeting antibiotics (e.g., β-lactams). These results highlight the importance of assessing the extracellular environment in determining antibiotic doses and the use of β-lactams antibiotics for treating infections caused by swarmer cells of *P. mirabilis* and *V. parahaemolyticus*.

## Introduction

Bacteria have evolved a variety of mechanisms to adapt to their physical environment. For example, in response to fluctuating environmental conditions, changes in biochemistry and gene regulation can alter bacterial shape and increase cell fitness. Cell filamentation is a commonly observed change in bacterial cell shape (1, 2) and has been described as a mechanism that enables bacteria to evade predation by the innate immune system during host infections (1).

In close proximity to surfaces, many bacteria alter their morphology and leverage cell-cell physical contact to move collectively to access new sources of nutrients and growth factors (3, 4). Referred to as ‘swarming’, this process is common among motile bacteria and has been connected to bacterial pathogenesis and infections (3, 4). Swarmer cells of *Salmonella enterica*, *Pseudomonas aeruginosa*, *Serratia marcescens*, and *Bacillus subtilis* have reduced antibiotic susceptibility—compared to vegetative cells—to a variety of drugs that alter protein translation, DNA transcription, and the bacterial cell membrane and cell wall (5–8). The specific biochemical and biophysical mechanisms underlying these observations are unknown.

Here, we describe physical changes in swarmer cells of the gram-negative pathogenic bacteria *Proteus mirabilis* and *Vibrio parahaemolyticus* that have the opposite effect: they increase the susceptibility of cells to cell wall-targeting clinical antibiotics. We found that large changes in the length of *P. mirabilis* and *V. parahaemolyticus* swarmer cells are accompanied by an increase in flexibility (i.e., a reduction in cell stiffness) that enables long cells to pack together tightly and form cell-cell interactions; maximizing cell-cell interactions promotes surface motility (9). Using biophysical, biochemical, and structural techniques, we quantified changes in the structure and composition of the *P. mirabilis* and *V. parahaemolyticus* cell wall in swarmer and vegetative cells and characterized their susceptibility to osmotic changes and cell wall-modifying antibiotics. Our results indicate that morphological changes that enable these bacteria to adapt to new physical environments come at a significant fitness cost, as cells become more susceptible to their chemical environment. In particular, changes in the composition and thickness of *P. mirabilis* and *V. parahaemolyticus* swarmer cells may make them more sensitive to osmotic changes and to cell-wall modifying antibiotics, thereby suggesting that these classes of drugs may be useful in treating infections of these bacteria (e.g., in urinary tract infections).

## Results

### The bending rigidity of *P. mirabilis* and *V. parahaemolyticus* cells decreases during swarming

During surface motility, *P. mirabilis* and *V. parahaemolyticus* cells grow into swarmers that are characteristically long (10-100 μm) and present a high surface density of flagella that enables them to translate through viscous environments (3, 10). We found that these swarmer cells display an unusual phenotype that is rarely observed among gram-negative bacteria: remarkable flexibility and a shape that is dynamically altered by adjacent cell motion and collisions (Fig. 1). The ability of *P. mirabilis* swarmer cells to maximize cell-cell contacts plays a role in their cooperative motility (10); our observations indicate that flexibility enables these long cells to optimize packing into multicellular structures that move cooperatively across surfaces.

**Fig. 1.**
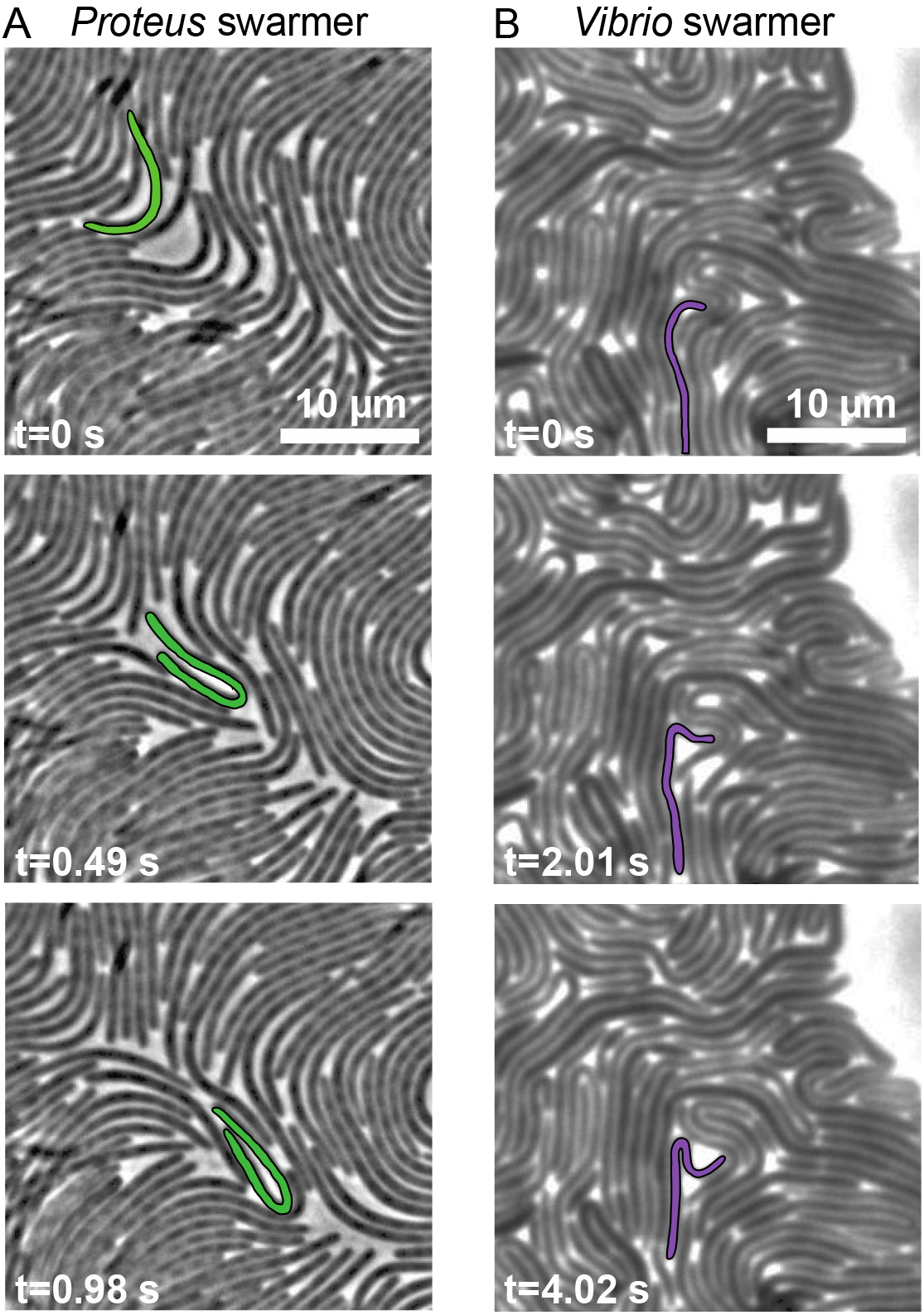
Images demonstrating the flexibility of *P. mirabilis* and *V. parahaemolyticus* swarmer cells. (*A*) Time-series of *P. mirabilis* swarmer cells in a colony actively moving across the surface of a 1.5% agarose gel. A representative cell, false colored green, has a generally straight shape at t=0 s and is bent in half at t=0.98 s. Most of the cells in this frame are bending during this imaging sequence. (*B*) A time-series of *V. parahaemolyticus* swarmer cells in a colony actively moving across the surface of a 1.4% agarose gel. A representative cell (false colored purple) has a generally straight shape at t=0 s and is bent in half at t=4.02 s.

Bacterial cell mechanics is generally attributed to the peptidoglycan (PG) layer of the cell wall, which has a thickness of ∼3-50 nm and surrounds the cytoplasmic membrane (11). Very little is known about mechanical regulation in bacteria (12–17) and we are unaware of studies connecting swarming to changes in cell mechanics. We quantified changes in swarmer-cell stiffness using cell-bending assays in a reloadable, poly(dimethylsiloxane) microfluidic system (Fig. 2 and S1) that is related to a method developed previously (18). In bending assays, we applied a shear fluid force to multiple filamentous cells, resulting in horizontal deflection of their cell tips (Fig. 2); fitting the deflection data to a mechanical model provided us with a value of the (flexural) bending rigidity of cells (Fig. S2). After a thorough comparison of several models of variable complexity, we found our results to provide a reasonable semi-quantitative estimate of bending rigidity. Introducing a reloadable mechanism enabled us to perform rapid bending measurements of *P. mirabilis* and *V. parahaemolyticus* swarmer cells after isolating them from swarm plates. Once removed from a surface, *P. mirabilis* and *V. parahaemolyticus* swarmer cells dedifferentiate, grow, and divide to form vegetative cells with a wild type length, requiring us to rapidly perform assays with swarmer cells after their isolation from surfaces. As a point of comparison, we filamented vegetative cells of *P. mirabilis* and *V. parahaemolyticus* using aztreonam—an inhibitor of the division specific transpeptidase PBP3—to match the length of swarmer cells (22.2 ±12.5 μm and 12.4 ± 8.2 μm respectively) and compared their bending rigidity values to swarmer cells. As a control, we measured the bending rigidity of cells of *Escherichia coli* strain MG1655 that we filamented using aztreonam, and determined the value to be 3.7 × 10^−20^ N m^2^ (Fig. 3); using a value for the thickness of the PG of 4-nm (19) yields a Young’s modulus of 23 MPa, which is close to values that have been reported previously and supports the choice of using aztreonam to filament cells, as it apparently has no effect on the bending rigidity of cells (12, 18). We assume that the effect of aztreonam on *P. mirabilis* and *V. parahaemolyticus* cells is similar to that which we measured for *E. coli*. We found a substantial decrease in the bending rigidity of swarmer cells of both *P. mirabilis* (∼26-fold) and *V. parahaemolyticus* (2.1-fold) compared to vegetative cells (Fig. 3), which is consistent with our observations of the flexibility of swarming cells by microscopy. *V. parahaemolyticus* vegetative cells were remarkably flexible: ∼154-fold more than *E. coli* cells and ∼58-fold more than *P. mirabilis* swarmer cells (Fig. 3).

**Fig. 2.**
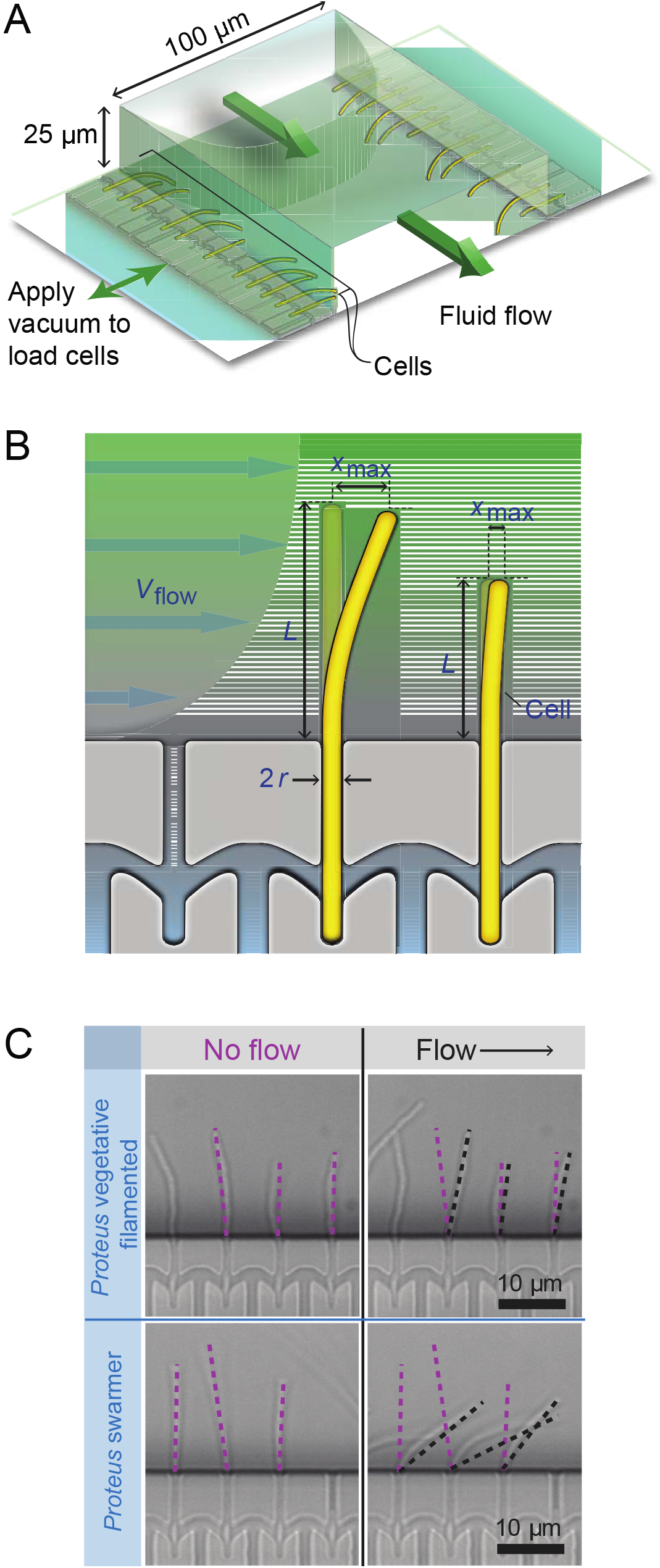
Using a reloadable microfluidic-based assay to determine bacterial cell stiffness. (*A*) Schematic of the microfluidic channel used to apply a user-defined shear force to bend filamentous or swarmer cells. Single-sided green arrows depict the flow of fluid through the central channel; the parabolic flow profile of the fluid is shown. Double-sided green arrows indicate the vacuum chamber used to load cells into side channels and to empty the device. (*B*) Cartoon of a flexible bacterium (left) and a stiff bacterium (right) under flow force (*V*_flow_). *x*_max_ indicates the deflection of cells in the flow; 2*r* = cell diameter; *L* = cell length in contact with the flow force. (*C*) Representative images of filamentous cells of *P. mirabilis* in no flow (left) and flow (right) conditions (top) and *P. mirabilis* swarmers (bottom). Purple dashed lines indicate the position of a cell tip under no flow conditions and black dashed lines illustrate the position after flow is applied using a gravity-fed mechanism. The arrow indicates the direction of fluid flow in the channel.

**Fig. 3.**
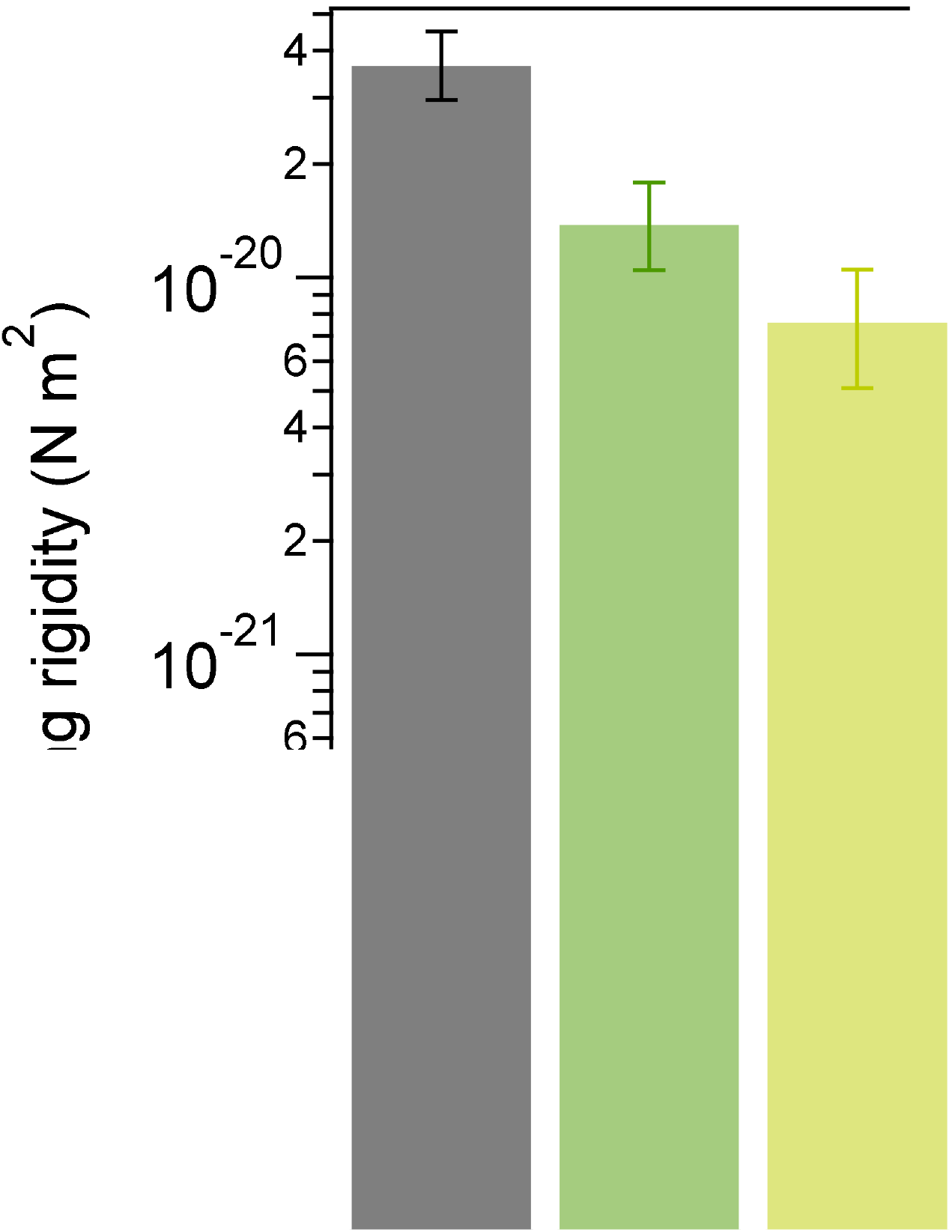
*P. mirabilis* and *V. parahaemolyticus* swarmer cells have a lower bending rigidity than vegetative cells. We measured the bending rigidity of *P. mirabilis* and *V. parahaemolyticus* swarmer cells and filamentous vegetative cells in a microfluidic flow assay and included measurements of vegetative *E. coli* cells. *P. mirabilis* swarmers exhibited 26-fold lower bending rigidity than vegetative cells; *V. parahaemolyticus* swarmers were 2-fold less rigid than vegetative cells. Overexpression of FlhDC (from the plasmid-encoded *pflhDC*) had little effect on the stiffness of *P. mirabilis* vegetative and swarmer cells. The values for each cell type are an average of two fitting models (see SI) and the brackets bars represent the upper and lower limits of the two models. *n* > 100 cells were used for each cell type from at least 3 independent experiments. The 95% confidence intervals associated with the fits are shown in Fig. S19. The plot has a logarithmic y-axis scale.

To confirm that using aztreonam to inhibit PBP3 and produce filamentous cells does not change the cross-linking density of PG at the division plane and alter cell mechanics, we compared bending rigidity values of cells treated with aztreonam and cells filamented by overexpressing SulA, a protein that prevents polymerization of the division protein FtsZ and blocks cell division. Both mechanisms of filamenting *E. coli* cells produced similar bending rigidity values: 3.8 × 10^−20^ N m^2^ (SulA) and 3.7 × 10^−20^ N m^2^ (aztreonam) (Fig. S3A).

During swarming, *P. mirabilis* and *V. parahaemolyticus* cells dramatically increase their density of flagella. Since flagella are attached to these cells through a protein complex that extends through the PG (referred to as the basal body), it is possible that these structures create local defects in the PG that alter its mechanical properties. To test whether the density of flagella on swarming cells is responsible for the changes in cell mechanics that we observe, we performed bending rigidity measurements (Fig. S3B) on two K-12-derived strains of *E. coli* that have substantially different flagella densities (Fig. S3C). Our results indicated no appreciable change in stiffness values (bending rigidity of 4.1 × 10^−20^ N m^2^ at high flagella density and 3.7 × 10^−20^ N m^2^ at low flagella density; Fig. S3B).

Overexpressing FlhDC—the heterohexameric activator that is important for swarming—in vegetative cells growing in liquid produces a phenotype that replicates many of the characteristics of swarmer cells, including increased cell length and flagella density (20). A relationship between FlhDC production and cell mechanics is untested. To test whether FlhDC is connected to changes in swarmer cell stiffness, we overexpressed the protein from the plasmid-encoded genes *pflhDC* in filamentous cells of *P. mirabilis* and measured their bending rigidity. We detected a ∼1.8-fold difference in bending rigidity between wildtype (1.4 × 10^−20^ N m^2^) and *pflhDC*-containing *P. mirabilis* vegetative cells (7.8 × 10^−21^ N m^2^), and approximately equal bending rigidity between wildtype swarmer and *pflhDC*-containing *P. mirabilis* swarmer cells (5.5 × 10^−22^ and 7.4 × 10^−22^ N m^2^, respectively) (Fig. 3). These results indicate that FlhDC overexpression during swarming is a minor contributor to the mechanical phenotype of swarming cells; the majority of the mechanical changes we observe arise from another regulatory pathway(s).

### *P. mirabilis* and *V. parahaemolyticus* swarmer cells are more sensitive to osmotic stress than vegetative cells

To complement cell stiffness measurements, we also studied changes in the mechanics of swarmer cells and filamented vegetative cells by measuring their changes in length and width in response to changes in osmotic pressure, which cause cells to shrink (in a NaCl solution) and swell (in H_2_O) (Fig. S4). We filamented *P. mirabilis*, *V. parahaemolyticus*, and control *E. coli* cells using aztreonam to create a range of cell lengths that matched the lengths of *P. mirabilis* and *V. parahaemolyticus* swarmer cells. We expected that low values of bending rigidity and compositional and structural changes in PG would cause swarmer cells to elongate in response to changes in osmotic pressure (in comparison to filamented, vegetative cells). We found that *P. mirabilis* and *V. parahaemolyticus* swarmer cell extension in response to osmotic shock was significantly larger than filamented, vegetative cells (Fig. 4). To circumvent the cellular production of osmolytes to protect cells from large changes in osmotic pressure during shock (typically produced within 1 min) (21), we used a microfluidic device to rapidly switch (<5 s) between H_2_O and a NaCl solution and measured changes in cell length before cells adapted (Fig. S4).

**Fig. 4.**
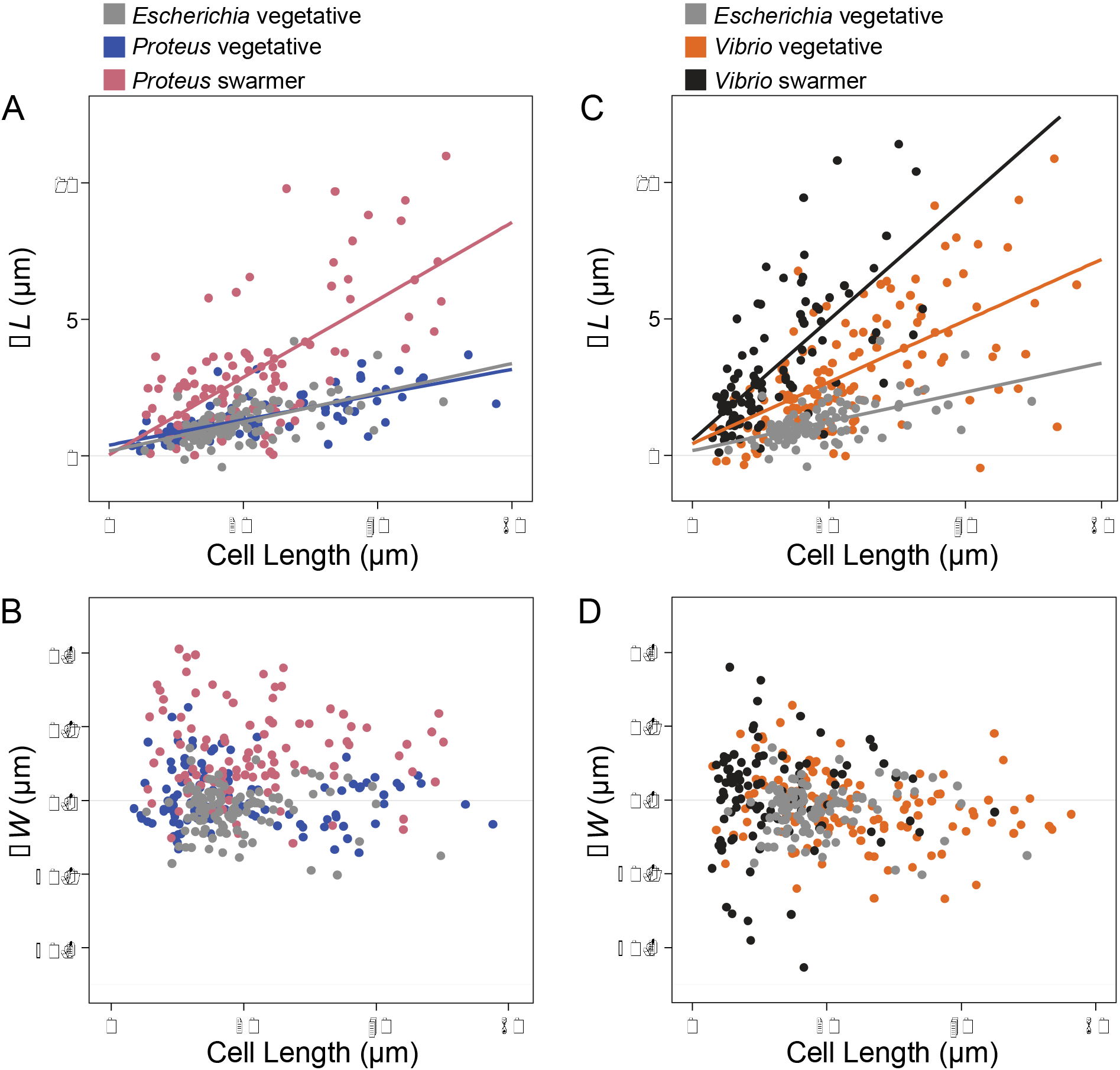
Swarmer cells increase in cell extension during osmotic shock. We calculated Δ*L* as (cell length in water – cell length in 1 M NaCl) and performed a similar calculation for Δ*W*, substituting cell width. Cell length indicates length prior to osmotic shock. We filamented all vegetative cells using aztreonam to grow them to lengths that were comparable with *P. mirabilis* and *V. parahaemolyticus* swarmers. Lines indicate linear fits to single-cell measurements (circles) of *n* > 100 cells from at least 3 independent experiments. (*A*) *P. mirabilis* swarmer cells have an increase in extension (Δ*L*) under osmotic shock compared to *E. coli* and *P. mirabilis* vegetative cells. (*B*) *V. parahaemolyticus* vegetative and swarmer cells have an increase in extension (Δ*L*) under osmotic shock compared to *E. coli*. (*C*) *P. mirabilis* swarmer cells have an increase in Δ*W* compared to *E. coli*; *P. mirabilis* vegetative cells display a slight decrease in width and increased cell length. (*D*) There was no observable change in Δ*W* of *V. parahaemolyticus* swarmer and vegetative cells.

Osmotic shifts produced similar changes in the length (Fig. 4A) and width (Fig. 4B) of vegetative filamented *P. mirabilis* cells compared to *E. coli* cells (Figs. 4A,B). In contrast, *V. parahaemolyticus* vegetative cells substantially increased in cell length (Fig. 4C) compared to *E. coli* cells, with no observable change in cell width (Fig. 4D). In response to osmotic upshifts (i.e., transitioning from H_2_O to NaCl), *P. mirabilis* and *V. parahaemolyticus* swarmer cells dramatically increased in cell length (Figs. 4A,C) (and to a lesser extent, cell width for *P. mirabilis* but not for *V. parahaemolyticus*; Figs. 4B,D) compared to vegetative cells. These results suggest that changes in *P. mirabilis* and *V. parahaemolyticus* swarmer cell stiffness make them more responsive to osmotic changes.

### Changes in PG composition of *P. mirabilis* and *V. parahaemolyticus* swarmer cells

We next studied the structural and biochemical mechanisms underlying the change in mechanical properties of swarmer cells. The connection between the structure of PG and cell mechanical properties—dating back to early antibiotic studies that revealed the penicillin binding proteins—suggested PG a starting point for these studies.

PG consists of the disaccharide building block β-(1,4)-N-acetylmuramic acid/N-acetyl-glucosamine (MurNAc-GlcNAc) in which a pentapeptide is attached to each 3’-OH group on MurNAc. Cross-linking between adjacent pentapeptides creates a mesh-like polymeric layer, and altering its structure and composition affects cell-mechanical properties (14, 15). To determine whether the PG composition of *P. mirabilis* and *V. parahaemolyticus* cells changes during swarming, we isolated PG sacculi from vegetative and swarmer cells and used ultra performance liquid chromatography-mass spectrometry (UPLC-MS) to quantify its chemical composition (Fig. S5). As the PG composition of *V. parahaemolyticus* has not yet been reported, we characterized its muropeptide stem peptide using UPLC-MS/MS (Fig. S6 and Table S1). Similar to *E. coli* (22) and *P. mirabilis* (23), *V. parahaemolyticus* has a PG structure that is conserved across other gram-negative bacteria, in which the peptide stem consists of L-Ala-D-Glu-*meso*-diaminopimelic acid (*meso*-DAP)-D-Ala-D-Ala (Fig. S6).

Compared to vegetative cells, *P. mirabilis* swarmer cells contained fewer monomers (MurNAc-GlcNAc), more dimers, and more anhydrous-containing saccharides, which are found at the terminating end of glycan polymers (Fig. 5A) (11). We detected no differences in the relative abundance of trimers between swarmer and vegetative cells of *P. mirabilis* (Fig. 5A). The increase in anhydrous-containing saccharides that we observed in *P. mirabilis* swarmer cells was correlated with a decrease in polysaccharide length (Fig. 5B). A similar increase in anhydrous-containing saccharides and decreased length of polysaccharides occurred in *V. parahaemolyticus* swarmer cells (Fig. 5A,B). We found no change in cross-linking density between vegetative and swarmer cells of either *P. mirabilis* or *V. parahaemolyticus* (Fig. 5C).

**Fig. 5.**
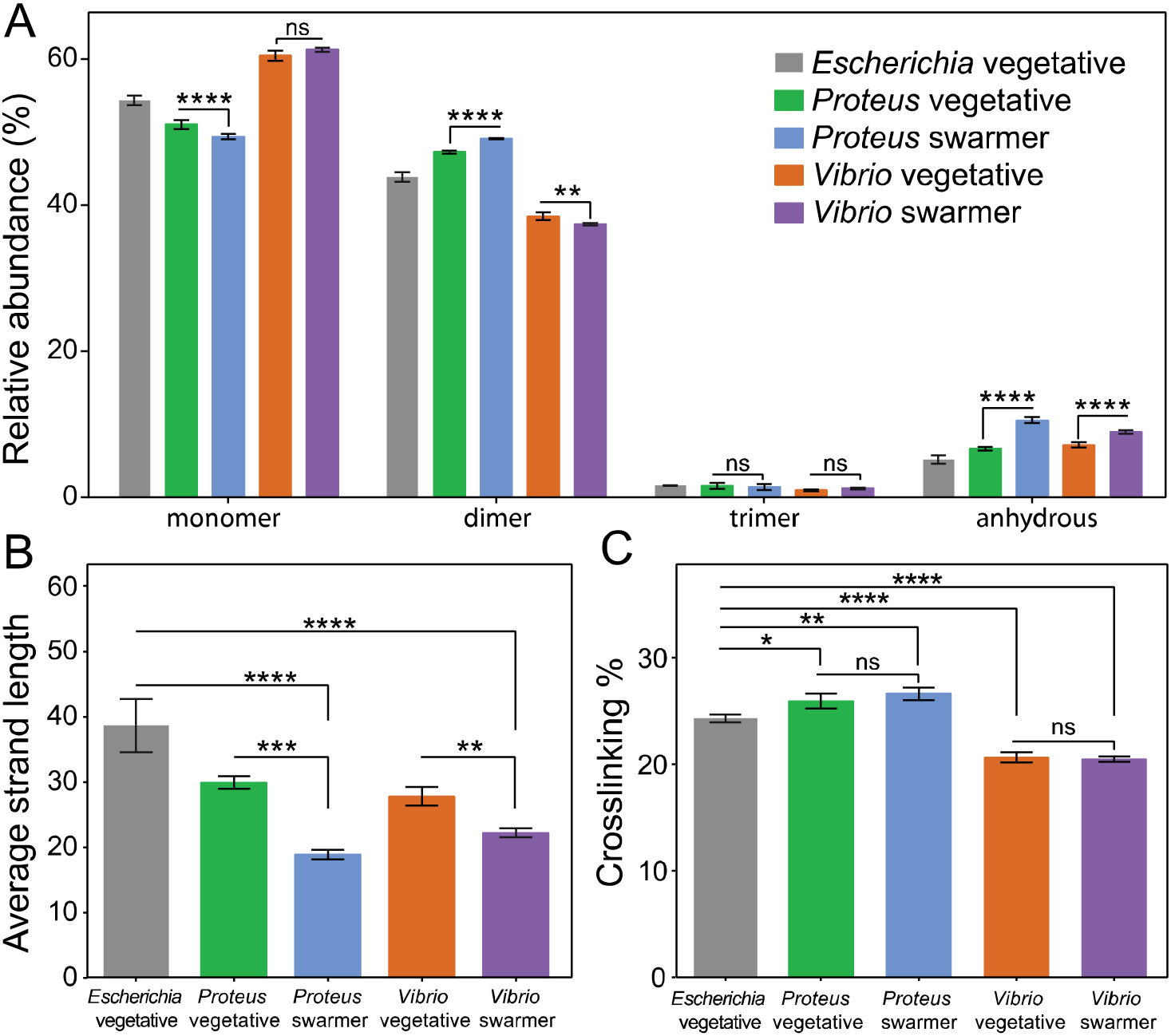
Alterations in the PG muropeptide composition of *P. mirabilis* and *V. parahaemolyticus* swarmer cells. (*A*) UPLC-MS data reveal that the muropeptide composition of *P. mirabilis* and *V. parahaemolyticus* vegetative and swarmer cells differs slightly in the abundance of monomer, dimer, and anhydrous-terminated saccharides. (*B*) We observed a relative increase in the amount of anhydrous-containing saccharides in swarmers consistent with a decrease in polysaccharide strand length. (C) There was no change in PG cross-linking of *P. mirabilis* and *V. parahaemolyticus* vegetative and swarmer cells, although *V. parahaemolyticus* does display a lower level of cross-linking. *n* = 3 biological replicates. Error bars represent the standard deviation of the mean. For (*A-C*), significance was determined via two-way analysis of variance: *P ≤ 0.05, **P ≤ 0.01, ***P ≤ 0.001, ****P ≤ 0.0001, ns = not significant (P > 0.05).

### Swarmer cells have a reduced PG thickness and display membrane defects

Changes in the thickness of the PG layer and structure of the cell envelope may also explain the observed decrease in swarmer cell stiffness. To identify changes in PG thickness of swarmer cells, we isolated intact sacculi from *P. mirabilis* vegetative and swarmer cells and measured the thickness of dried sacculi using tapping-mode atomic force microscopy (AFM) (Fig. 6A). Differences in the nanoscopic appearance of the sacculi of different cells were not observed by AFM (Fig. S7). The thickness of isolated, dry *P. mirabilis* swarmer cell sacculi (1.0 ± 0.2 nm) was reduced ∼1.5-fold compared to vegetative cells (1.5 ± 0.2 nm) (Fig. 6A). *V. parahaemolyticus* swarmer cells (0.6 ± 0.1 nm) exhibited a similar ∼1.2-fold decrease in thickness compared to vegetative cells (0.8 ± 0.2 nm). Earlier AFM measurements of isolated sacculi indicated that dehydration reduced the thickness of *Escherichia coli* PG by ∼2x, which we used to estimate the dimensions of hydrated PG from *P. mirabilis* (3.1 and 2.1 nm for vegetative and swarmer cells, respectively) and *V. parahaemolyticus* (1.7 and 1.4 nm, respectively) (24). A comparison of PG thickness and cell bending rigidity suggested that the relationship between these data is approximately exponential (R^2^=0.9874) (Fig. 6B).

**Fig. 6.**
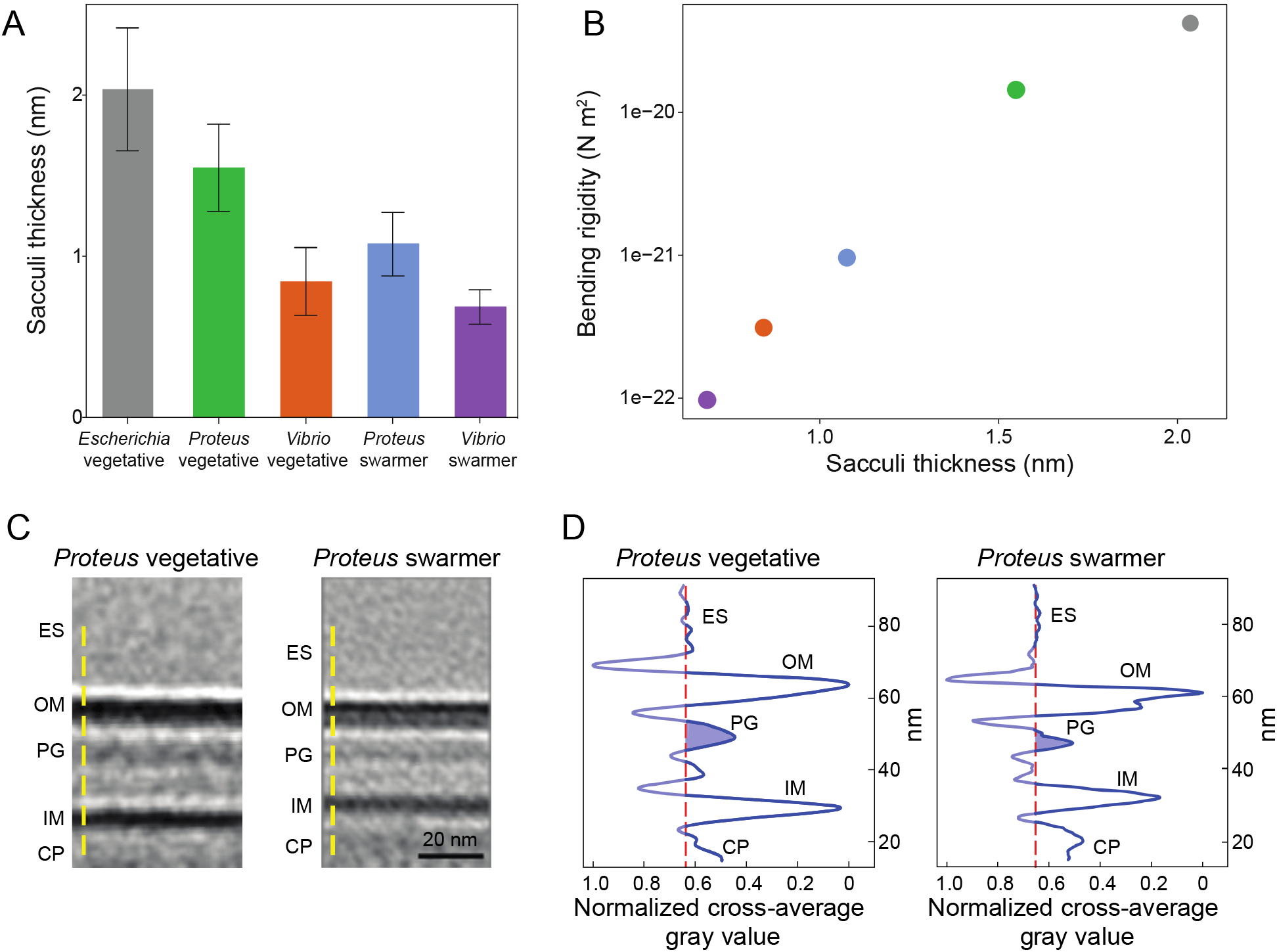
AFM reveals the PG layer of *P. mirabilis* and *V. parahaemolyticus* swarmer cells is thinner than in vegetative cells and ECT demonstrates a reduced membrane-to-membrane distance. (*A*) Sacculi were isolated from cells, dried, and imaged by AFM. The thickness of swarmer cell PG was reduced compared to vegetative cells. We analyzed >65 vegetative cells of *E. coli*, *P. mirabilis*, and *V. parahaemolyticus*, >65 *P. mirabilis* swarmer cells, and 7 *V. parahaemolyticus* swarmer cells. Error bars represent the standard deviation of the mean. (*B*) Bending rigidity and cell wall thickness display an approximately exponential relationship (R^2^=0.9874). (*C*) Sub-tomogram-averaged ECT volume images of the *P. mirabilis* vegetative (left) and swarmer (right) cell wall. Two central slices of sub-tomogram average volume images with normalized image densities are shown. Yellow dashed line indicates the orientation used for gray-value measurements. ES, extracellular space; OM, outer membrane; PG, peptidoglycan; IM, inner membrane; CP, cytoplasm. (*D*) The density profile of sub-tomogram-averaged ECT volume images reveals a reduced membrane-to-membrane distance in swarmer cells. The vertical axis is the normalized gray value with the darkest value equal to 0 and the lightest value equal to 1. The red dashed line denotes the average gray value of the extracellular space and serves as a reference for the background; the blue shaded area indicates the thickness of the putative PG layer.

A caveat with conversions between dried and hydrated values is that they are most accurate for PG that best mimics the structure and composition (e.g., crosslinking and polysaccharide composition) of the reference material: *E. coli* PG. Alterations in the polysaccharide length of PG from *P. mirabilis* and *V. parahaemolyticus* may be more elastic and stretch out during drying, thereby appearing to have a thickness that is reduced compared to *E. coli* PG. Our control measurements on isolated, dry *E. coli* sacculi yielded a thickness of 2.0 nm, which varies from the value of 3.0 nm published by Yao et al. using the same technique (24). A difference of ∼30% between these *E. coli* measurements may arise for several reasons, including: variations in physical conditions that impact AFM measurements, improvement in the resolution of AFMs, and/or the precision of fitting force curves.

The relatively low variability in the values we measured for isolated sacculi from *P. mirabilis* and *V. parahaemolyticus* cells (∼16-25%) using AFM demonstrated a consistent reduction in PG thickness for swarmer cells, suggesting that our measurements are sufficient for comparing PG from vegetative and swarming cells, and demonstrating a connection between PG and changes in cell mechanics. In contrast, the variability in AFM measurements and analysis makes us less comfortable comparing absolute values of PG thickness in our studies to those reported for bacteria in other papers.

To complement AFM measurements, we attempted to measure the thickness of native PG using electron cryotomography (ECT) on intact vegetative and swarmer cells (Figs. 6C,D, S8, and S9). Although we were unable to resolve the thickness of native PG by direct ECT measurements, we noticed that the sub-tomogram-averaged ECT volumes of the *P. mirabilis* cell wall (Fig. 6C) indicated the distance between the inner and outer membranes of swarmer cells was smaller than in vegetative cells (Figs. 6C,D). *P. mirabilis* vegetative cells had a characteristically smooth membrane (Fig. S8A,B) that was similar to the presentation of the membrane found along the lateral, cylindrical walls of swarmer cells (Fig. S8C,D). In contrast, the polar regions of both *P. mirabilis* and *V. parahaemolyticus* swarmer cells had an undulating outer membrane suggestive of an altered structure (Fig. S8E,F and S9D-F), and *V. parahaemolyticus* cells had significant defects in their cell-envelope, including, membrane budding, vesicle formation, and ruptured cell walls (Fig. S9D-F).

### Increased susceptibility of *P. mirabilis* and *V. parahaemolyticus* swarmers to the cell wall-targeting antibiotics

Although swarming colonies of bacteria display resistance to many antibiotics (5), our experiments suggest *P. mirabilis* and *V. parahaemolyticus* swarmer cells have changes in PG structure, composition, and properties that may increase their susceptibility to cell wall-targeting antibiotics. To determine the sensitivity of *P. mirabilis* and *V. parahaemolyticus* swarmers to the cell wall-targeting antibiotics cephalexin (inhibits PBP3 and 4) (25) and penicillin G (inhibits PBP3, 4, and 6), we measured swarmer single-cell growth in the presence of the minimum inhibitory concentration (MIC) of antibiotics in a microfluidic device. We determined MICs of cephalexin against vegetative cells of *E. coli* MG1655 (1X MIC = 6.25 μg/mL, 32X MIC = 200 μg/mL), *P. mirabilis* (100 μg/mL), and *V. parahaemolyticus* (50 μg/mL). The MIC of penicillin G against vegetative cells of *P. mirabilis* was 12.5 μg/mL. We were unable to measure MICs for these antibiotics against swarmer cells because they dedifferentiate at shorter time scales than the measurements used in the MIC protocol. Instead, we chose 3 h of incubation for our measurement for three key reasons: 1) swarmer cells do not dedifferentiate over this time period; 2) a past study demonstrated that treating cells with cephalexin for 3 h was sufficient to kill ∼100% of a population of *E. coli* cells (26); and 3) ∼2 h of cephalexin treatment was sufficient to observe cell lysis of *E. coli* cells measured by single cell growth (27).

We constructed a simple microfluidic flow cell to maintain the swarming phenotype of cells during experiments and supply cells with a source of continuously replenished fresh growth media to ensure exponential cell growth and a constant concentration of antibiotics. Channel walls consisted of the biocompatible polymer, poly(dimethylsiloxane) (PDMS). Cells were in contact with the channel floor, which consisted of a layer of a 2% agarose gel. See Figure S12 for a cross-section of the device. We found that the survival of *P. mirabilis* (66%) and *V. parahaemolyticus* (64%) vegetative cells treated with 1X MIC of cephalexin was slightly higher than *E. coli* cells (55%) (Figs. 7A, S10). Treating *P. mirabilis* and *V. parahaemolyticus* swarmer cells with 1X MIC of cephalexin reduced survival to 37% and 19%, respectively (Figs. 7A, S10), indicating their increased susceptibility. Rates of cell survival in the presence of penicillin G were similar to the use of cephelexin (Fig. 7A). We characterized dead cells using microscopy to measure membrane blebbing, cell lysis, and changes in the refractive index of cells.

**Fig. 7.**
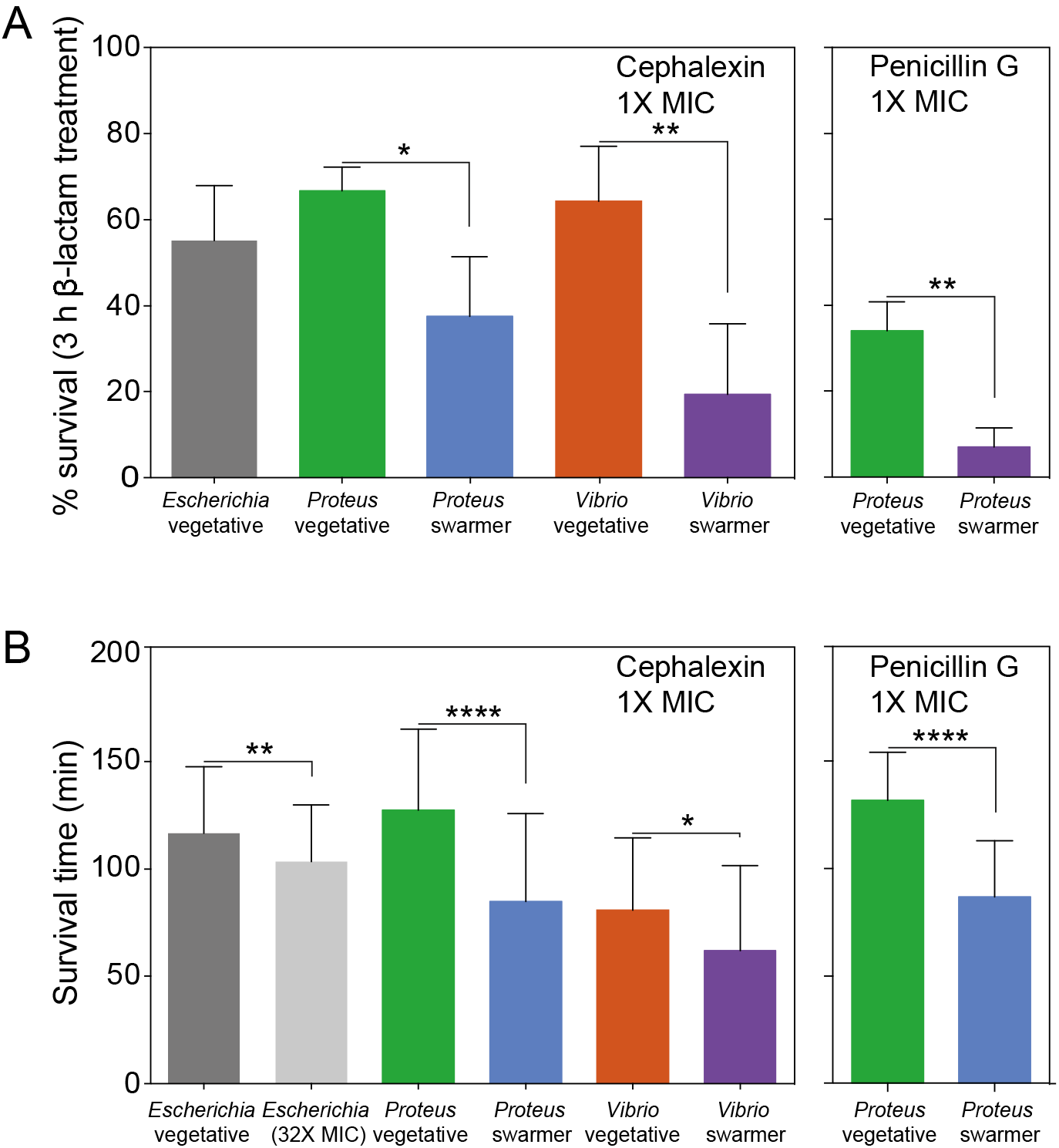
Swarmer cells are more susceptible to antibiotics that target the cell wall than vegetative cells. (*A*) Survival of cells treated with 1X MIC of cephalexin after 3 h of incubation. We define percent survival as (cell count_no lysis_ / cell count_total_) × 100. *P. mirabilis* and *V. parahaemolyticus* swarmer cells exhibit decreased levels (∼30%) of survival compared to vegetative cells; *n* ≥ 90 cells from at least two independent experiments. A similar decrease occurred when *P. mirabilis* was treated with penicillin G; *n* ≥ 77 cells from at least two independent experiments). Error bars represent the standard deviation of the mean. (*B*) After exposure to cephalexin or penicillin G, the survival time of *P. mirabilis* and *V. parahaemolyticus* swarmers was 2∼3 fold lower than for vegetative cells. Survival time was determined for ≥ 49 cells that lysed from at least 2 independent experiments. Significance in *A-B* was determined two-tailed t-test: *P ≤ 0.05, **P ≤ 0.01, ****P ≤ 0.0001.

Cell wall-targeting antibiotics are most effective against actively growing cells (28, 29) and therefore increases in growth rate will reduce cell survival. To determine whether a growth rate phenotype explains our observations, we normalized data for cell length, and were unable to detect a change in the growth rate of swarmer cells when they were treated with these two antibiotics (Fig. S11). We measured the mean survival time of cells (the amount of time elapsed after treatment with drugs before cell death) for *P. mirabilis* and *V. parahaemolyticus* swarmer cells treated with cephalexin and penicillin G (Fig. 7B) and found that the survival time for these swarmer cells was lower than that for vegetative cells (Fig. 7B), which is consistent with alterations in the cell wall.

## Discussion

*P. mirabilis* is commonly associated with complicated urinary tract infections and increased mortality in cases of bacteremia (30). Swarming is hypothesized to enable *P. mirabilis* cell movement from the ureter to the kidney; in support of this hypothesis, swarming deficient mutants have lower rates of host infection (31). We found that during swarming, *P. mirabilis* and *V. parahaemolyticus* cells become more flexible due to changes in the thickness and composition of the PG. Although we were unable to precisely measure the thickness of isolated PG, the decrease in the distance between the inner and outer membranes of swarmers may reflect a reduction in PG thickness. Several recent studies suggest how changes in the cell wall influence cell stiffness. A recent study revealed the connection between changes in the biochemical regulation of D-Ala-D-Ala and *Pseudomonas aeruginosa* cell stiffness in cells in which this biochemical machinery is altered (32). This current study and the *P. aeruginosa* paper (32) demonstrate that it is still difficult to quantitatively understand the scaling between the magnitude of the biochemical changes in cells and changes in cell stiffness. The range of techniques now available for quantitatively measuring single cell stiffness will enable the development of this relationship.

We found that *P. mirabilis* and *V. parahaemolyticus* swarmer cells were more susceptible to large changes in cell length than vegetative cells during osmotic shift experiments. Although these changes are consistent with what would be expected from a reduction in PG thickness in swarming cells, they are also relevant to other cell wall changes. We also observed that the length of polysaccharides was reduced in swarming cells, which would not obviously lead to large changes in cell elongation during osmotic shifts if the organization of PG in these cells follows what is known about the structure of PG in *E. coli* cells (33): polysaccharides reported to be arranged circumferentially around cells with the peptide cross-links oriented parallel to the long axis of cells. Instead, we expect that a decrease in polysaccharide length makes swarmer cells more prone to changes in diameter. *P. mirabilis* swarmer cells appear to increase in diameter during osmotic shifts (with no change occurring for vegetative cells), however the data are noisy and not accurately fit with standard models; we were unable to observe a trend with *V. parahaemolyticus* swarmer cells (Fig. 5).

Another recent study demonstrated the role of the outer membrane in the cell stiffness of gram-negative bacteria (34). Chemical and genetic changes in the outer membrane increased the deformation of the cell envelope to stretching, bending, and indentation forces and increased the amount of cell lysis arising during osmotic shock. It is possible that the morphological changes in swarmer cell membranes that we observed during ECT experiments indicate membrane perturbations that contribute significantly to changes in cell stiffness. Swarming in *Proteus* and *Vibrio* is well known to occur in parallel with the secretion of lipopolysaccharides (3, 4), the source of which is presumably the outer membrane of these bacteria, which may alter cell stiffness.

This paper demonstrates for the first time the ability of bacteria to modulate their cell stiffness (beyond L-forms). Presumably this phenotype (i.e., more flexible) conveys an adaptive advantage for cells that is supported by observations from many labs that the motility of communities of swarmer cells is enhanced when cell-cell contacts are increased. The adaptive advantage of swarming in *P. mirabilis* and *V. parahaemolyticus* however appears to be offset by a decrease in their fitness, as cells become more sensitive to osmotic changes and cell wall-targeting antibiotics, thereby creating an “Achilles heel” for targeting this phenotype in infectious diseases. This study highlights the plasticity of bacteria (1), the continuing need for methods of evaluating chemotherapies that measure efficacy against cells in a physiological state relevant to specific infections, and the reconsideration of cell wall-targeting antibiotics for treating UTIs and other infections in which *P. mirabilis* and *V. parahaemolyticus* may be present.

## Materials and Methods

### Bacterial strains and cell culture

*P. mirabilis* strain HI4320, *V. parahaemolyticus* LM5674, *Escherichia coli* MG1655 (CGSC #6300), and plasmids *pflhDC* (10) and *psulA*(18) were used for experiments used in this paper. *P. mirabilis* was grown in PLB nutrient medium consisting of 1% peptone (weight/volume), 0.5% yeast extract (w/v), and 1% NaCl (w/v). *V. parahaemolyticus* was grown in nutrient medium (HI medium) consisting of 2.5% heart infusion (w/v) and 2.5% NaCl (w/v). *E. coli* was grown in lysogeny broth (LB) consisting of 1% tryptone (w/v), 0.5% yeast extract (w/v), and 1% NaCl (w/v). All strains were grown at 30 °C with shaking at 200 rpm.

### Immunostaining flagella on *E. coli* cells to determine flagella surface density

As a control to determine the influence of the surface density of bacterial flagella on measurements of bending rigidity, we imaged two *E. coli* MG1655 strains with different flagella densities using a polyclonal flagellin antibody and an immunostaining procedure for flagella visualization (35).

### Calculating vegetative and swarmer cell division times

We prepared vegetative cells by diluting an overnight culture 1:200 in fresh medium and grew the cells at 30 °C with shaking at 200 rpm to an optical density (OD; λ=600 nm) of 0.6. We prepared swarmer cells as described previously (35). We monitored the growth of *P. mirabilis*, *V. parahaemolyticus*, and *E. coli* vegetative cells over 120 min at 30 °C in the microfluidic flow chamber device described in the next section. We performed a similar experiment with *P. mirabilis* and *V. parahaemolyticus* swarmers growing over 65 min at 30 °C to determine the amount of time elapsed before cells divided. We collected images of cells at 1-min intervals and determined the division time for a maximum of 10 generations (for vegetative cells) and 4 division events (for swarmer cells).

### Measuring the sensitivity of cells to osmotic shock in a microfluidic device

We prepared filamentous vegetative cells of *P. mirabilis* and *V. parahaemolyticus* by diluting an overnight culture of cells 1:200 into fresh medium. We then grew cultures at 30 °C with shaking for 1 h, added aztreonam (MP Biomedicals) to a final concentration of 10 μg/mL and grew cells for an additional 70 min. Swarmer cells were prepared as described previously (35). To maximize the number of cells attached to a surface in the microfluidic device (flow chamber construction described below), we concentrated cells to an OD_600_ of 8 in 100 μL of liquid nutrient medium.

We prepared the microfluidic device for experiments by flowing 10 μL of undiluted Cell-Tak (Corning) into the device and incubated it for 10 min at 25 °C. Next, we flowed 20 μL of a suspension of cells (OD_600_=8) through the device, then repeated with another 20 μL aliquot of cell suspension. To aid the adherence of the highly motile swarmer cells to the Cell-Tak-coated surface, we centrifuged the device for 5 min at 300 × *g* in a centrifuge (Beckman Coulter) equipped with a swinging bucket rotor.

For osmotic shock experiments, we filled separate 1-mL syringes (BD) with a solution of UltraPure distilled water (Invitrogen) or 1 M of a NaCl solution prepared in ddH_2_O and connected the syringes using a three-way valve. We mounted the microfluidic device on a TE2000-E inverted microscope and imaged cells in the chamber at 25 °C (decreasing the temperature reduced growth and cell division). Prior to osmotic shock, we flowed 200 μL of fresh liquid media (PLB for *P. mirabilis*, LB for *E. coli*, and HI for *V. parahaemolyticus*) through the device to remove cells that were not adhered to the channel surface. While imaging, we flowed a 1 M NaCl solution through the channel and observed cell plasmolysis, immediately after which we flowed ddH_2_O through the channel and observed cells elongating. We collected images of hundreds of single cells after exposing them to three conditions: 1) isotonic culture (PLB, LB, HI); 2) hypertonic shock (1 M NaCl); and 3) hypotonic shock (ddH_2_O).

To determine changes in cell size and width, we extracted individual cell contours from the images taken for each treatment using MATLAB 2014a (MathWorks) and used MicrobeTracker (36) to determine cell lengths (*L*) and widths (*W*). From these measurements we calculated Δ*L* (*L*_hypotonic_ − *L*_hypertonic_) and Δ*W* (*W*_hypotonic_ − *W*_hypertonic_) for vegetative and swarmer cells of *P. mirabilis* and *V. parahaemolyticus*.

### Determining the minimum inhibitory concentration (MIC) of vegetative cells of *P. mirabilis* and *V. parahaemolyticus*

We used the micro-dilution protocol (37) to determine the MIC of cephalexin in accordance with Clinical and Laboratory Standards Institute. Briefly, we added 400 μg/mL of cephalexin (BP Biomedicals) or penicillin G (BP Biomedicals) to the first well of a 96-well microplate (Nunc) and diluted these antibiotics 2-fold across adjacent wells (wells #1-11); well 12 was a no-drug control. We determined the MIC after 16 h of growth at 30 °C with shaking by identifying the lowest concentration of cephalexin and penicillin G that inhibited cell growth by visual inspection. The MIC was determined from three replicate plates. We determined the MIC of cephalexin against *E. coli* MG1655 (1X MIC = 6.25 μg/mL, 32X MIC = 200 μg/mL), *P. mirabilis* (100 μg/mL), and *V. parahaemolyticus* (50 μg/mL). The MIC of penicillin G against *P. mirabilis* was 12.5 μg/mL.

### Antibiotic treatment of vegetative and swarmer cells and measurement of their growth in a microfluidic device

We prepared vegetative and swarmer cells of *P. mirabilis* and *V. parahaemolyticus* as described above. Prior to use, we diluted cells 1:100 in fresh medium to a cell density that enabled us to image many individual cells simultaneously in the microfluidic device described in the section above.

To prepare the microfluidic device for monitoring growth, we applied a 250 μm-thick layer of PDMS prepolymer to the surface of (#1.5, 35 × 50 mm) cover glass (FisherBrand) using a spincoater (Laurell Technologies) and polymerized the polymer overnight at 60 °C. We subsequently removed a 6 mm × 4 mm rectangle of PDMS from the center of the cover glass using a scalpel. We applied transparent tape to both sides of this rectangular well and pipetted 100 μL of media containing a 2% (w/v) solution UltraPure agarose (Invitrogen) into the well. We placed a (#1.5, 22 × 30 mm) cover glass (FisherBrand) on top of the liquid agarose (to flatten the agarose surface), pressed the cover glass against the transparent tape, and solidified the agarose at 25 °C. Next, we removed the #1.5 cover glass, tape, and any residual agarose from the PDMS surface. We pipetted 2 μL of a suspension of cells on the agarose pad surface, waited until the excess liquid had evaporated or been absorbed by the agarose, carefully removed the agarose layer, inverted it, and placed it back into the well such that the cells were positioned between the agarose surface and the cover glass. We placed the PDMS flow chamber used in the construction of the osmotic shock microfluidic device on the PDMS-coated cover glass and aligned it such that the agarose pad was centered in the flow channel (Fig. S12).

For antibiotic treatment of cells in the microfluidic cell growth device, we filled a 6-mL syringe (Norm-Ject) with cephalexin or penicillin G at a concentration corresponding to 1X MIC dissolved in nutrient medium. We supplied a constant flow of 20 μL/min to the device using a syringe pump (Harvard Apparatus) and monitored the growth of individual cells during antibiotic exposure using a TE2000-E inverted microscope. The stage and objective heater were maintained at 30 °C. Images were collected every 1 min for 3 h.

Due to the aberrant shape of cells treated with cephalexin and penicillin G (Fig. S10), we were unable to use an automated script (e.g., MicrobeTracker) to determine cell death. Instead, we visually determined the time of death for individual cells when they exhibited three phenotypes that collectively indicate cell death: blebbing (membrane swelling), lysis (bleb rupture), and disappearance of cells in phase contrast microscopy (loss of cytoplasmic material).

### Determining cell growth rates in the presence of antibiotics

We prepared vegetative and swarmer cells of *P. mirabilis* and *V. parahaemolyticus* as described above. We monitored individual cell growth at 30 °C in the presence of 1X MIC of cephalexin and pencillin G in our microfluidic growth device (described above) by collecting an image every 1 min for 15 min. To calculate the growth rate, we first extracted individual cell contours at each time point using MicrobeTracker and determined cell length. To take into account the differences in starting length between *P. mirabilis* and *V. parahaemolyticus* cells, we normalized the change in length to the initial cell length (Δ*L/L*_0_) at each time point. To determine the growth rate for individual cells, we fit their relative length over time to an exponential function using GraphPad Prism 6.0 (GraphPad Software).

### Measuring cell envelope architecture and peptidoglycan thickness using electron cryotomography

We prepared vegetative and swarmer cells of *P. mirabilis* and *V. parahaemolyticus* and concentrated them to an OD_600_=10. For ECT, we mixed vegetative and swarmer cells with bovine serum albumin-treated 10-nm diameter gold particles that served as fiducial markers, applied them to electron microscopy grids, and plunge-froze them in a mixture of liquid ethane and propane, as described previously (38). Grids were stored in liquid nitrogen until imaging.

We acquired images on a 300 KeV Polara transmission electron microscope (FEI) with a GIF energy filter (Gatan) and a K2 Summit direct detector (Gatan). We collected tilt series from −60° to +60° with 1° increments using UCSFtomo software with a defocus of −10 μm and a total dosage of 190 e^−^/Å^2^ (39) at a magnification of 27500×. Tomograms were calculated using IMOD software (40).

For sub-tomogram averaging, smooth and flat membrane regions were chosen by eye; a volume of 40 × 70 × 12 voxels (62 nm × 109 nm × 19 nm) was centered using the outer membrane and extracted. We aligned 38 extracted “membrane fragments” from four *P. mirabilis* vegetative cells and 42 fragments from nine *P. mirabilis* swarmer cells and averaged them in PEET (41). The densities from two averaged membranes were scaled to match each other using IMOD (40), cross-averaged density profiles were measured using ImageJ 1.50c (42), and figures were generated in OriginPro (OriginLab).

### Determining the thickness of *P. mirabilis* and *V. parahaemolyticus* sacculi (isolated peptidoglycan) by atomic force microscopy (AFM) in ambient conditions

We isolated *P. mirabilis* and *V. parahaemolyticus* swarmer cells, concentrated them at 800 × *g* for 10 min, removed the supernatant, flash froze the cell pellet in liquid nitrogen, and stored it at −80 °C. Swarmer cell pellets were thawed at 4 °C and pooled for isolation of sacculi (intact peptidoglycan). To increase the efficiency of cell lysis prior to isolating sacculi, we resuspended vegetative and swarmer cell pellets in 3 mL of cold 1X phosphate-buffered saline (ThermoScientific), then lysed cells with a tip sonicator (Qsonica) for ∼10 s at a power setting of 75%. We confirmed cell lysis using optical microscopy. We isolated sonicated cells, resuspended sacculi in 20 (*V. parahaemolyticus* swarmers) or 200 μL of ddH_2_O (all other cells), immediately flash froze the sacculi in liquid nitrogen, and stored them at −80 °C.

To prepare sacculi for AFM, we transferred 10 μL of sacculi thawed at 4 °C to a new microcentrifuge tube placed in a bath sonicator (Branson) that was cooled with ice for 10 min to aid in the dispersal of sacculi without affecting sacculus architecture (43). After sonication, we pipetted 10 μL of the sacculi onto freshly cleaved mica (Ted Pella), dried the mica under nitrogen gas, washed it 3x with 1 mL of ddH_2_O (filtered through a 0.2 μm-diameter pore filter (Corning)), dried the sacculi under nitrogen gas, and imaged immediately after preparation.

We performed AFM using a Catalyst AFM (Bruker) operating in tapping mode in ambient conditions (air) with an aluminum reflex-coated silicon AFM probe (Ted Pella; k = 40 N/m). Before imaging, AFM probes were auto-tuned using Nanoscope 8.15 (Bruker). We collected all images at high resolution (512 × 512 pixels) with a scan speed of 1 Hz and analyzed images using NanoScope Analysis 1.4 (Bruker). Prior to determining sacculi thickness, we flattened (0^th^ order) all images to remove variations in surface thickness. Thickness was determined perpendicular to the long axis; we avoided surface debris, folds in the sacculi, and trapped material in sacculi (Fig. S7).

### Determining the composition of peptidoglycan in *P. mirabilis* and *V. parahaemolyticus* vegetative and swarmer cells using ultra high performance liquid chromatography/mass spectrometry (UPLC/MS)

We prepared vegetative and swarmer cells of *P. mirabilis* and *V. parahaemolyticus* as described above. To purify peptidoglycan for UPLC/MS, we carried out a previously reported isolation technique for Gram-negative bacteria (44) with the following modifications. Briefly, after trypsin (Sigma) inactivation, we incubated the sacculi in 1 M HCl solution (Fluka) for 4 h at 37 °C to remove any O-acetylation from the peptidoglycan, which is present in *P. mirabilis* (45). Then, the sacculi were washed three times in ddH_2_O, resuspended in 500 mM boric acid (Sigma) [pH 9] to OD_600_=3, and mixed with 1/10 the volume of mutanolysin (Sigma). The sample was incubated 16 h at 37 °C with 200 rpm shaking. The next day, the samples were centrifuged for 10 min at 9500 × *g*, pelleting the remaining insoluble material. The supernatant was removed and put into a glass vial. To reduce the isolated muropeptide fragments, we added 50 μL of 20 μg/mL sodium borohydride (Sigma) in 500 mM boric acid [pH 9] and incubated the mixture for 30 min at 25 °C. We adjusted the pH of the solution to 2-3 by adding 50% phosphoric acid (Fluka), then filtered the muropeptide solution through a Duropore polyvinylidine fluoride filter (0.22-μm pores; Millex) into a clean vial. Vials were immediately stored at −80 °C until use within 1 week of muropeptide isolation.

For UPLC/MS, we injected 7.5 μL of purified muropeptides on a Cortecs 2.1 × 100 mm C18 column (Waters) packed with 1.6 μM-diameter particles and equipped with a Cortecs C18 guard column (Waters). The column temperature was maintained at 52 °C using an Acquity standard flow UPLC system equipped with an inline photodiode array (Waters). For muropeptide separation by UPLC, we used solvent A (Optima LCMS-grade water with 0.05% trifluoroacetic acid) and solvent B (30% (v/v) Optima LCMS-grade methanol in Optima LCMS-grade water with 0.05% trifluoroacetic acid) (Fisher Scientific). Muropeptides were eluted from the column with a gradient of increasing solvent B (1 min hold at 1% B, ramp to 99% B over 59 min, hold at 99% B for 5 min, then decrease to 1% B over 1.5 min, then hold at 1% B for 4.5 min) at a flow rate of 0.2 mL/min. We analyzed the eluent from the column using a Bruker MaXis Ultra-High Resolution time-of-flight 4G mass spectrometer (Bruker Daltonic) with either an MS method or a data-dependent, top 3 MS/MS method. For both methods, capillary voltage was set to 4100 V, the nebulizer pressure was 2.0 bar, and the drying gas was set to 6.0 L/m at 220 °C. Muropeptides were detected at λ=205 nm by MS. We determined the peptidoglycan composition of *E. coli*, *P. mirabilis*, and *V. parahaemolyticus* vegetative and swarmer cells by comparing MS/MS fragmentation patterns using DataAnalysis version 4.2 (Bruker) (Fig. S6). Muropeptides were identified according to mass values using DataAnalysis 4.2. We calculated muropeptide masses using ChemDraw 14.0 (CambridgeSoft) (Table S1). We quantified the corresponding UV (λ=205) absorbing peaks (Fig. S5) identified by MS (Bruker) from which we calculated peptidoglycan cross-linking density and strand length (46) for each species. Statistical significance was determined using GraphPad Prism 6.0

### Fabrication of microfluidic devices for measuring cell bending and cell growth

Masters for the cell bending device and the cell growth/osmotic shock device were fabricated on separate 3” silicon wafers using SU8 photoresist that was exposed on a Heidelberg μPG101 mask writer (Heidelberg Instruments, Heidelberg, Germany) and developed. The bending device (Fig. S1) is a 2-layer device. The first SU8 layer consists of ∼1 μm tall channels for capturing cells. The wafer was then coated with the second SU8 layer (∼25 μm tall) that formed a central flow channel. The flow chamber (Fig. S12), consists of a single channel with a length of 10 mm, a width of 5 mm, and a height of 50 μm. The volume of the flow chamber is ∼10 μL. After developing the masters for both devices, we used the masters to emboss layers of poly(dimethylsiloxane) (PDMS), punched holes for inlets and outlets, cleaned the surfaces with tape. If the device was attached to a glass coverslip it was treated with oxygen plasma and immediately sealed against a plasma-treated # 1.5 cover glass (24 × 50 mm Fisherbrand) to form a permanent seal.

### Measuring the bending rigidity of *P. mirabilis*, *V. paramaemolyticus*, and *E. coli* cells

We used streak velocimetry to determine the profile of fluid flow rates in the central channel of the microfluidic device driven by gravity flow. We added fluorescently labeled 0.22 μm microspheres (Polysciences) diluted ∼1:10000 in ddH_2_O containing 0.01% Brij-35 (Sigma) to the microfluidic channel and collected videos of the fluorescent beads moving through the channel at focal planes 2 μm apart. We analyzed the movies with custom-written code in Igor Pro 6.37 (WaveMetrics). Briefly, we applied a Gaussian blur and threshold to each frame and used the thresholded image to establish a region that was fit to a function based on a 2D Gaussian. We used the image exposure time and length of the streaks—taking into account the size of the microspheres—to calculate microsphere velocity. We mapped the velocity profile within the channel by analyzing several hundred microspheres. We binned the velocity data into a 3D matrix and fit it to a Poiseuille function, letting the velocity coefficient, height, and width float. The calculated velocity coefficient was used as an input to the cell bending fitting function.

The system for gravity flow pumping (Fig. S13) consisted of two 6-mL syringes (Norm-Ject), one 60-mL syringe (BD), one 1-mL syringe (Norm-Ject), a ruler, and a VC-6 channel valve controller (Warner Instruments) connected to a VC-8 mini-valve system (Warner Instruments) that drives a three-way solenoid valve (The Lee Company). Syringe #1 (6-mL) was mounted to an immobile post, positioned 40 cm above the table surface; this syringe was used to load cells. Syringe #2 (6-mL) was positioned 75-cm above the table surface, was connected to a stage that could be raised and lowered vertically, and was used to apply flow force in the device. Syringe #3 (60-mL) was mounted to an immobile post, positioned 75-cm above the surface of the table, and was used to apply an opposing flow force at the device outlet and to collect spent media/cells. We attached two-way valves, a blunt-end needle, and tubing to each syringe, including syringe #4 (1-mL). We joined the two 6-mL syringes using a Y-junction connector that led to the VC-8-mini-valve system inlet. The outlet of the valve system was connected to the microfluidic system. A ruler on an immobile post indicated the ‘0 position’ (no pressure).

For cell bending experiments, we prepared filamentous vegetative cells and swarmer cells of *P. mirabilis* and *V. parahaemolyticus*; cells were normalized to an optical density (OD; λ=600 nm) of 1. Prior to starting a measurement, syringes 1 and 2 were flushed with ddH_2_O and fresh medium. Syringe 2 was filled with 4 mL of medium, syringe 3 was filled with 30 mL of ddH_2_O, and syringe 4 was filled with 0.8 mL of ddH_2_O. We added cells to syringe 1 and flowed them through the tubing until they reached the device outlet. To load the cells into the side channels of the device (from the central channel), we applied a suction force using the 1-mL syringe. After the side channels were loaded with cells, we adjusted the height of syringe 2 until no flow occurred in the device. Syringe 2 was then raised 7 cm from the no-flow position (a height of 0 cm). We collected images of cells in the channel when syringe 2 was at positions 0 cm and 7 cm using a Zeiss Axiovert 100 inverted microscope (Zeiss) equipped with an iXon3 CCD (Andor), a 63X Plan-APOCHROMAT oil objective (Zeiss), and Micro-Manager 1.4.16. After collecting cell-bending deflections for all loaded cells, we expelled cells from the side channels, flowed liquid through the device for 15 s, and reloaded the side channels with new cells.

We analyzed images to determine cell deflection under flow using custom image-analysis software written in Igor Pro 6.37. The cell-bending model (Supplementary Information) is a differential equation that lacks an analytical solution and thus requires calculation of a numerical solution. To determine the bending rigidity of cells, we wrote custom fitting code in Igor Pro 6.37 that uses a variety of input parameters, including channel dimensions, fluid velocity, bending rigidity of the cell, cell radius, and cell length. The function numerically calculates the maximum deflection based on our model. This function is integrated into a fitting algorithm to find a least-squares solution to a dataset of maximum deflections versus cell lengths with bending rigidity as the fitting parameter.

Our model for bending a cell under flow is based on the mechanics model of a suspended rod or cantilever bending under its own weight (see DERIVATIONS in Supplementary Information). Our experimental system was similar to that of Amir et al. (47). Although laminar flow is perpendicular to the long axis of cells in this system, the lateral deformations of cells that we measured were substantially larger (in our case, as large as 10 μm; compared to < 1 μm in Amir et al. (47). For this reason, many of the assumptions of the model presented by Amir et al. are not valid for our dataset. The model we developed to extract the bending rigidity of the bacterium takes into account the shape of the laminar flow profile, the angle of the cell against the flow profile, and the arc length of the cell (longer cells tend to fold over and do not penetrate as deeply into the flow profile). Additional data and mathematical derivations are described in Supplementary Information.

## Acknowledgements

We thank Linda McCarter for *V. parahaemolyticus* LM5674, Suckjoon Jun for plasmid *psulA*, Cameron Scarlett and Molly Pellitteri-Hahn for mass spectrometry support, and Julie Last for technical assistance with AFM measurements. This research was supported by the Bill and Melinda Gates Foundation (grant OPP1068092), NIH grant 1DP2OD008735-01, National Science Foundation grant DMR-1121288, a Mao Wisconsin Distinguished Graduate Fellowship (to M.R.), an NSF postdoctoral fellowship (#1202622 to P.M.O), and the Howard Hughes Medical Institute.

