## Supplemental Information for "Bacterial swarming reduces *Proteus mirabilis* and *Vibrio parahaemolyticus* cell stiffness and increases β-lactam susceptibility"

**for**

**Figures and Legends**

**Figure S1. Reloadable microfluidic device design.** A cartoon depicting the entire microfluidic device including inlet, outlet, vacuum inlet, and the central bending chamber. Inset cartoon of the loading channels depicts the central bending chamber in which cells are loading into the 1^st^ tier of channels using negative pressure. Inset cartoon of pillars (vacuum chamber): pillars are located in the vacuum chamber to ensure that the chamber does not collapse when the vacuum is applied. Green highlights the 1^st^ layer of device (1-μm thick); black highlights the 2^nd^ layer of device (25-μm thick).

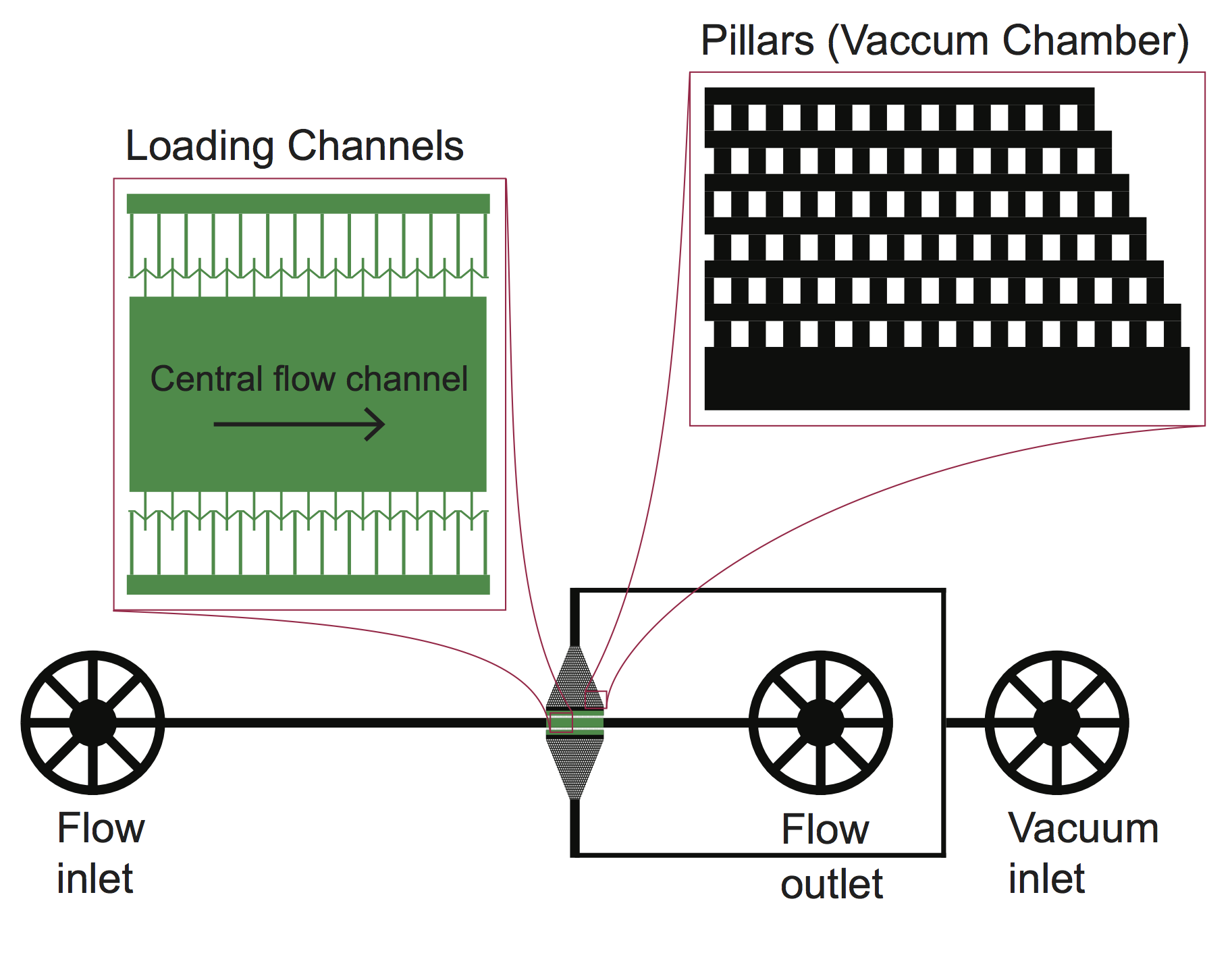

**Fig. S1**

**Figure S2. Deflection of vegetative filamented and swarmer cells immediately after flow-induced bending in microfluidic device.** Circles represent the deflection value of individual cells under fluid flow. Larger deflections (Figure S2C, E) indicate a decrease in cell stiffness. The black line represents a fit to two models (C1 and C2, see SI methods, below) to the data to determine the flexural rigidity (*n* > 200 cells, from at least 2 independent experiments).

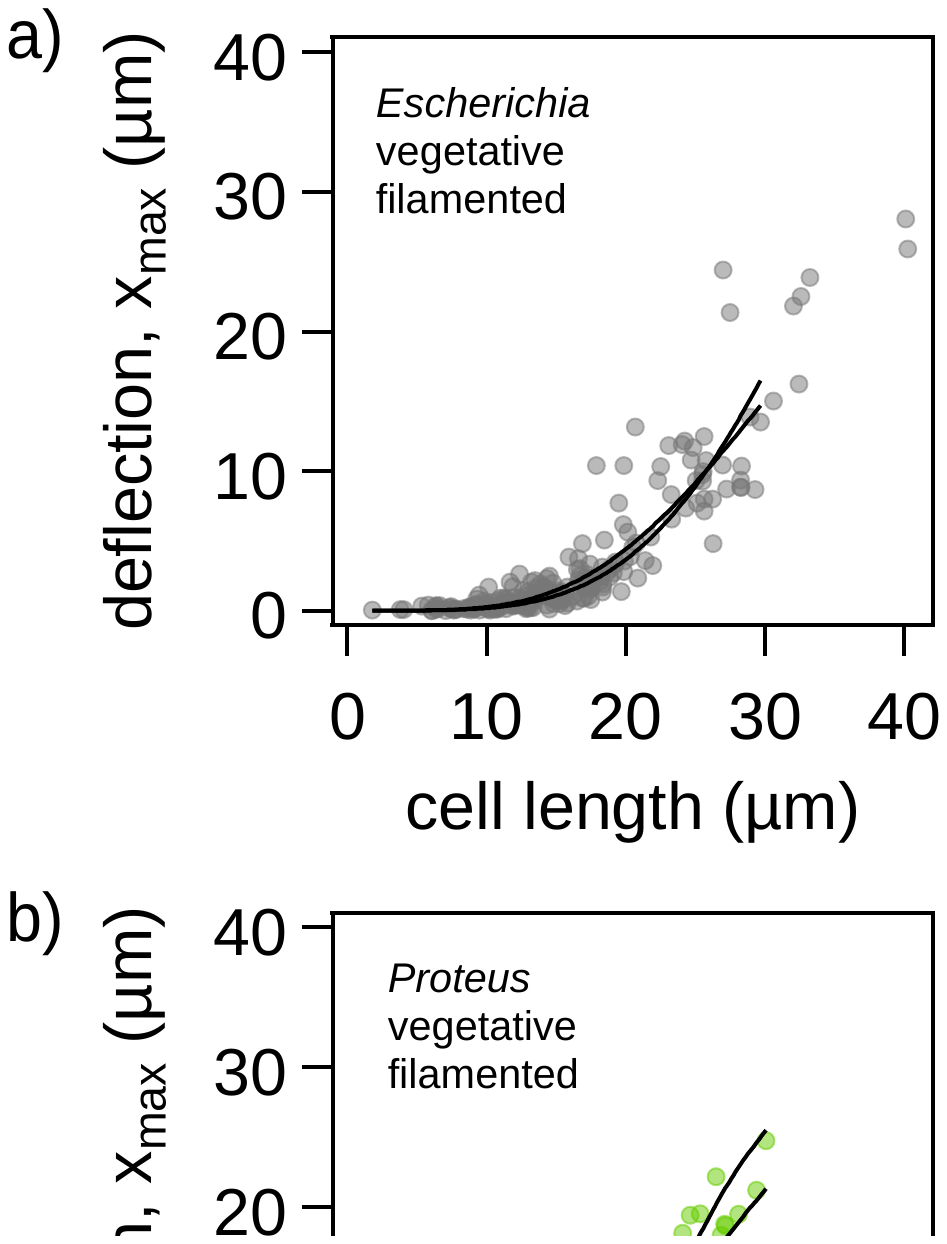

**Fig. S2**

**Figure S3. Flexural rigidity of *E. coli* is unaffected by treatment aztreonam or by an increase in flagella density.**

A) Bending rigidity is not affected by filamentation with aztreonam. Flexural rigidity of wild-type *E. coli* MG1655 cells filamented by expression of *SulA* (green) or treatment with aztreonam (gray). Flexural rigidity of cells was determined by fitting deflection data (Methods) from microfluidic-based bending assays. The data indicates the average of two computational models and the brackets represent the limits of these models. See ‘Derivations’ section for more information. (*n* > 100 cells).
B) Bending rigidity is not affected by an increase in number of flagella on the cell body. Data indicate the flexural rigidity of cells filamented with aztreonam: wild-type *E. coli* MG1655 (gray) and isolated *E.* coli MG1655 with an increased flagella density (red) (see Fig S3C). The data indicates the average of two computational models and the brackets represent the limits of these models. See ‘Derivations’ section for more information. (*n* > 100 cells).
C) Immunofluorescence images of wild-type *E. coli* MG1655 (top panel) and *E. coli* MG1655 cells with increased flagella density (bottom panel) filamented with aztreonam. Flagella were labeled with anti-FliC primary antibody and an Alexa Fluor 488 conjugated secondary antibody. Scale bar = 10 μm.

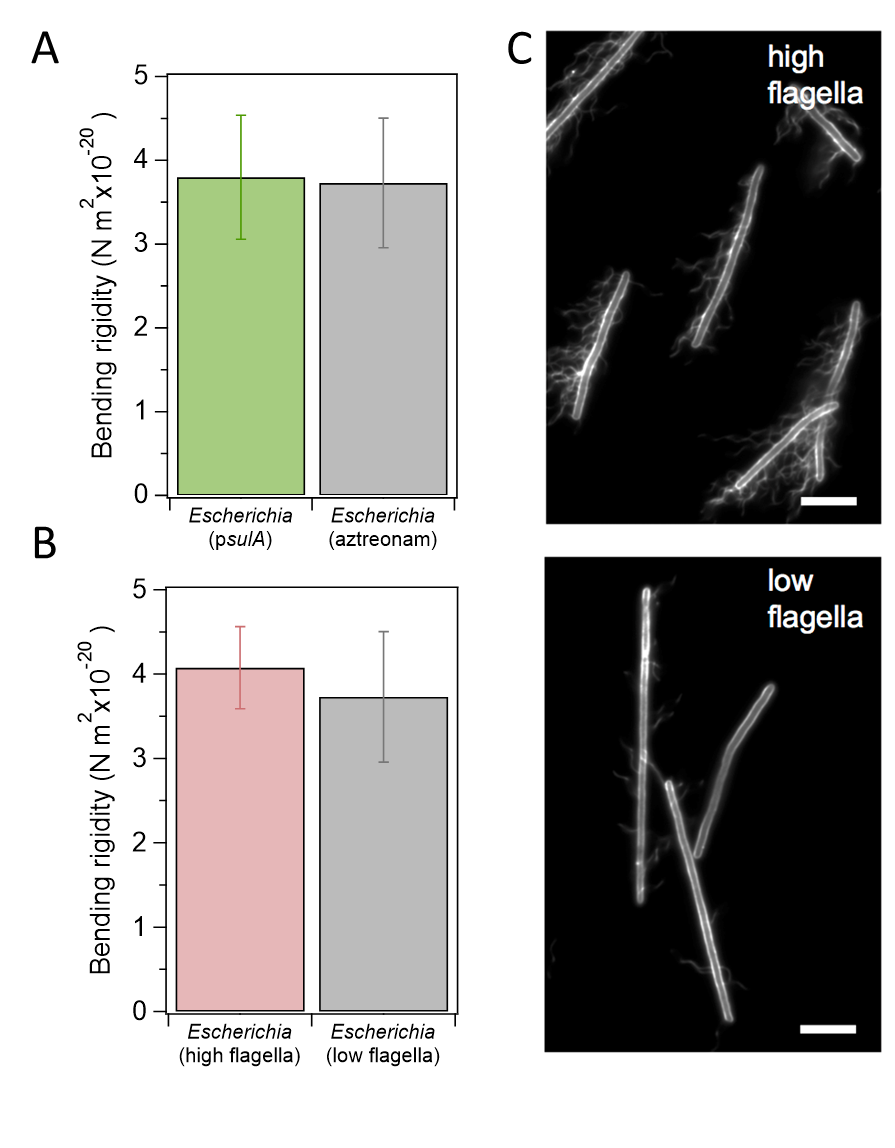

**Fig. S3**

### **Figure S4. Elongation of vegetative filamented and swarmer cells during osmotic shock.** Cells were attached to the glass surface using Cell-Tak. Prior to performing osmotic experiments, we flowed fresh media through the device to remove any non-adhering cells from the surface. We flowed a hypertonic solution (1M NaCl) solution through the device until we saw visible cell plasmolysis. Immediately after, we flowed the hypotonic solution (ddH_2_O) through the device until cells fully elongated. (A) Vegetative filamented cells and swarmer cells exposed to hypotonic (ddH_2_O) and hypertonic (1M NaCl) conditions to alter cell length through increasing and decreasing turgor pressure, respectively. Scale bar = 10 μm. (B) A cartoon depicting the response of cell length during osmotic; arrows indicate the direction of turgor pressure in the cell.

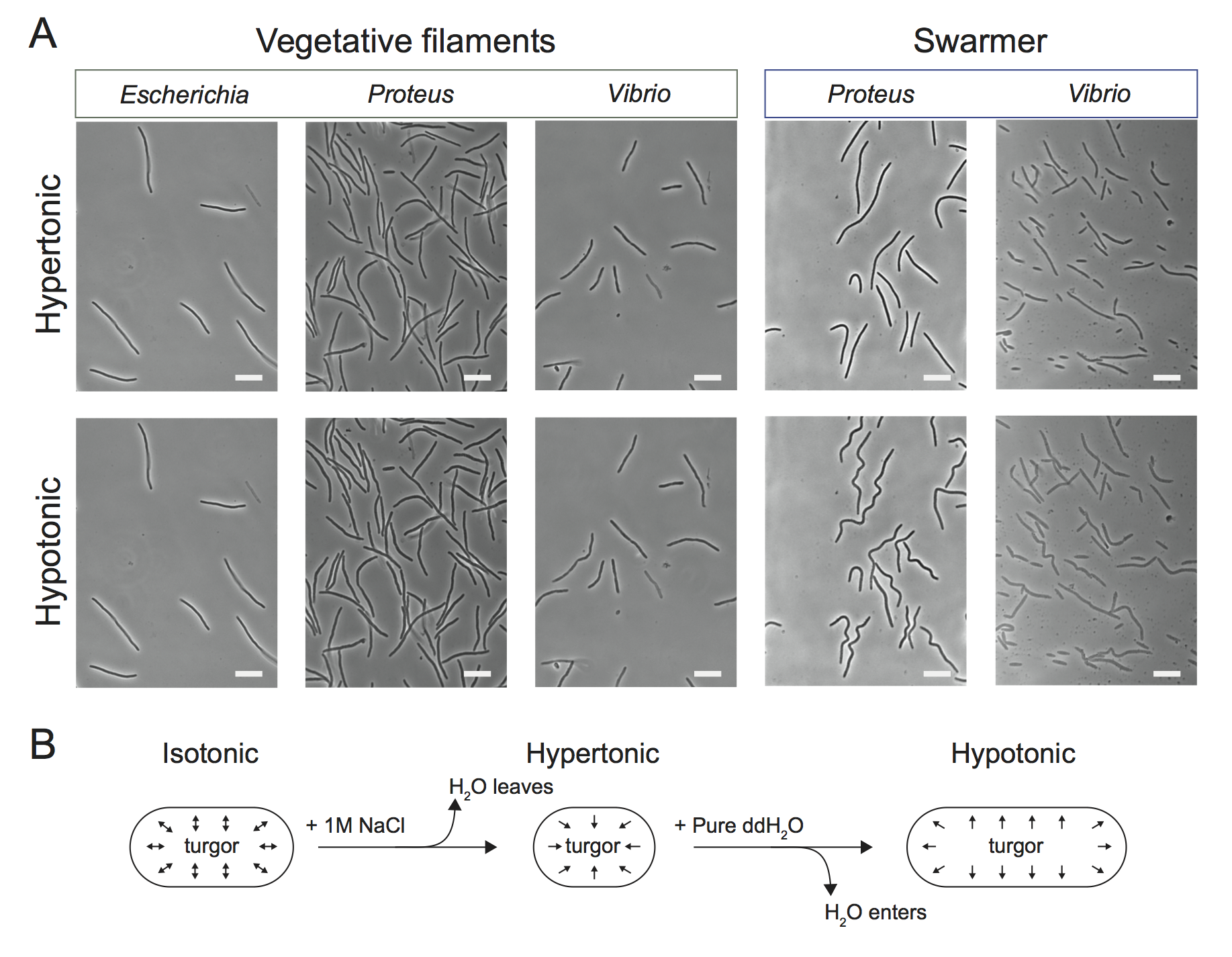

**Fig. S4**

**Figure S5. UPLC-MS data of muropeptides isolated from vegetative and swarmer cells.** Chromatograms of purified cell wall from *E. coli, P. mirabilis, V. parahaemolyticus* vegetative cells; and *P. mirabilis*, *V. parahaemolyticus* swarmer cells. We purified sacculi using the 24 h isolation method, digested them with mutanolysin, and analyzed them by UPLC-MS. Muropeptides were separated by UPLC using solvent A (water with 0.05% TFA) and solvent B [30% (v/v) methanol in water with 0.05% TFA]. Muropeptides were eluted from the column with a gradient of increasing solvent B at a flow rate of 0.2 mL/min; our gradient consisted of the following steps: hold 1 min at 1% B, ramp to 99% B over 60 min and hold at 99% B for 5 min; to prepare for the next sample we ramped back down to 1% B over 0.5 min and held at 1% B for 4.5 min. Table S1 lists the peaks that we identified and Figure 5 displays the quantification of peaks.

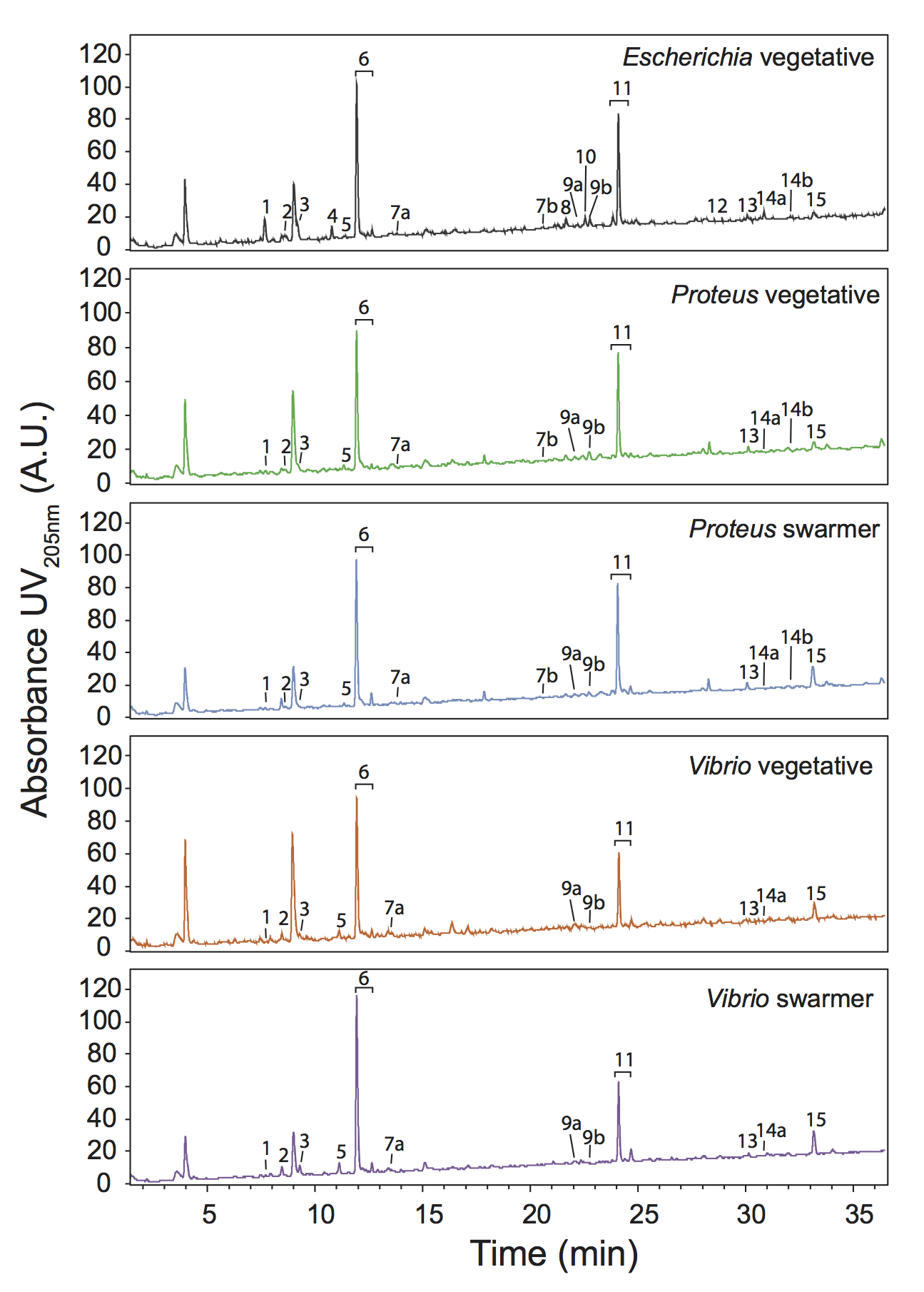

**Fig. S5**

**Figure S6. Determination of the muropeptide stem by MS/MS.** We analyzed the tetrapeptide peak (observed - 942.4036 m/z, calculated – 942.4155 m/z) by performing MS/MS on the parent ion to determine its amino acid composition. MS/MS confirmed that the *E. coli*, *P. mirabilis*, and *V. parahaemolyticus* muropeptide compositions are identical and consist of: L-Ala, D-Glu, meso-diaminopimelic acid, and D-Ala.

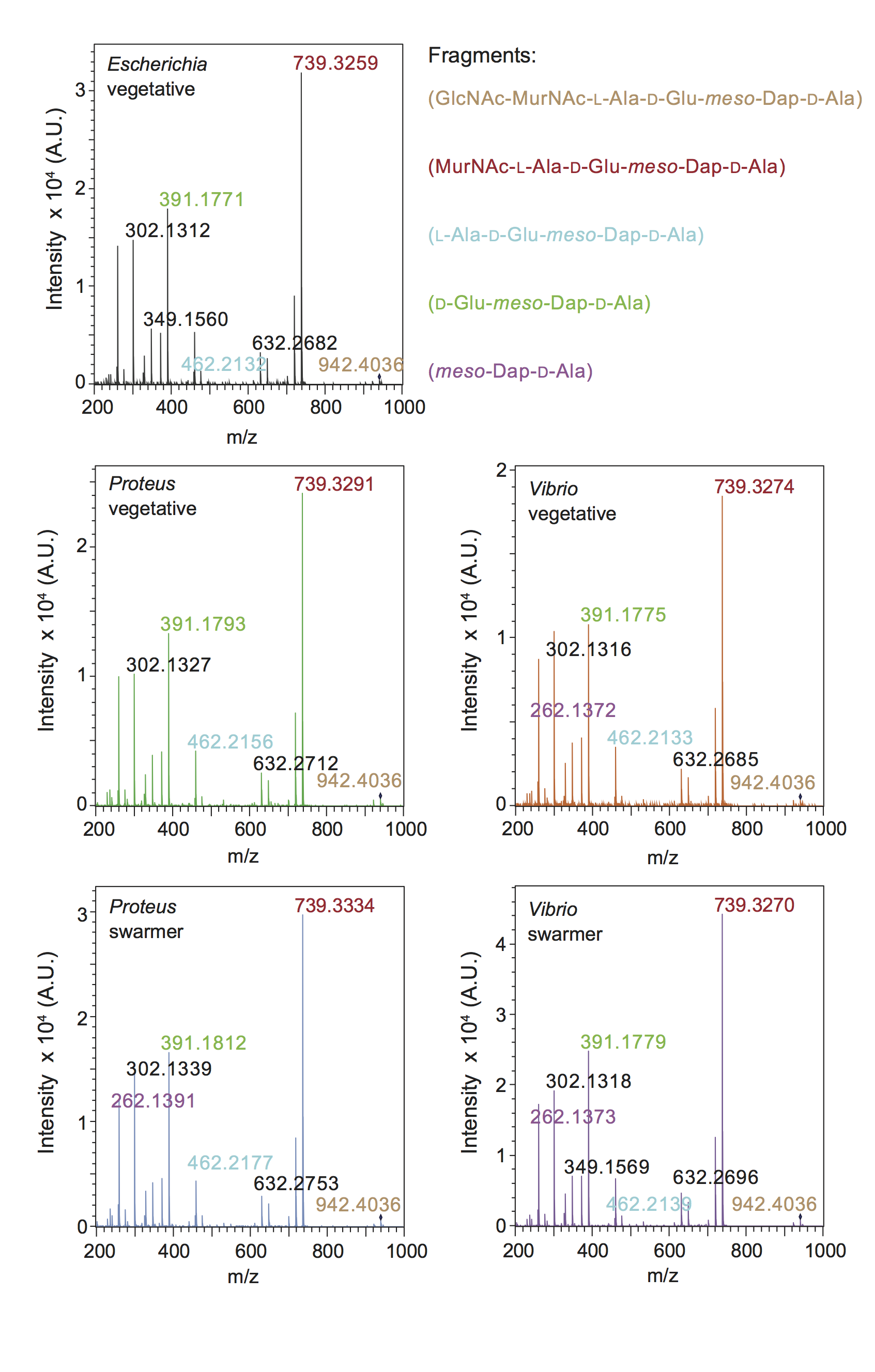

**Fig. S6**

**Figure S7. AFM images of isolated sacculi from *E. coli*, *P. mirabilis*, and *V. parahaemolyticus*.** A sample of dispersed sacculi was pipetted on freshly cleaved mica, dried under nitrogen gas, and imaged immediately. Imaging was performed using tapping mode AFM in ambient conditions (air). Images were collected at high resolution (512 x 512 pixels) with scan speed of 1 Hz. A) *E. coli* vegetative, B) *P. mirabilis* vegetative, C) *P. mirabilis* swarmer, D) *V. parahaemolyticus* vegetative, E) *V. parahaemolyticus* swarmer. Scale bar = 1 μm

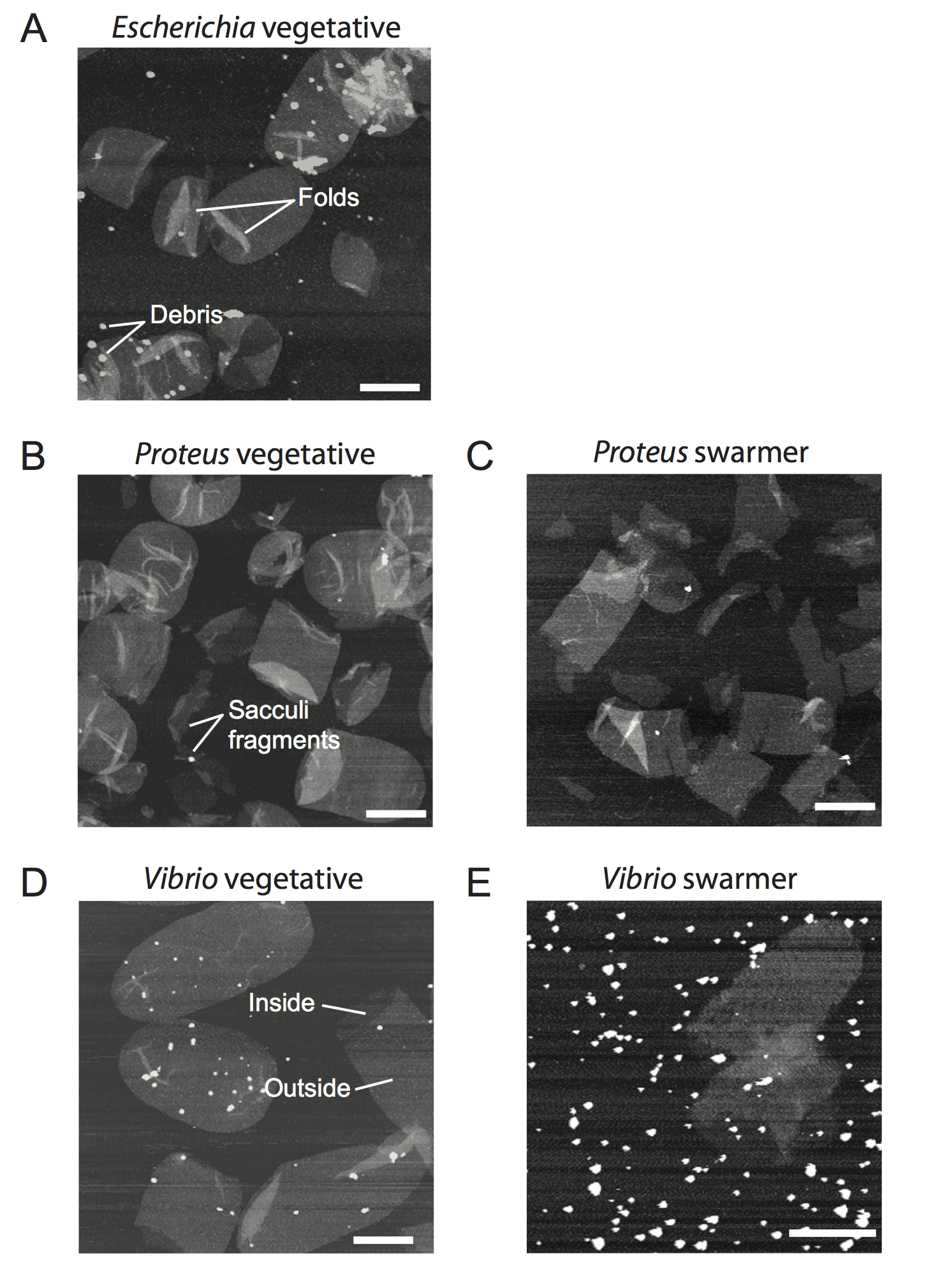

**Fig. S7**

**Figure S8. Electron cryotomography** **of *P. mirablis* vegetative and swarmer cells reveals decreased membrane stability.** *P. mirabilis* vegetative cells (A, B) have a smooth outer membrane and an increased distance between the inner and outer membrane (compared to swarmer cells). Swarmer cells (C-F) display a smooth lateral wall membrane (C, D) and a ruffled outer membrane near the pole (E, F) indicating a decrease in membrane stability. Representative features are labeled in figure panels A-F.

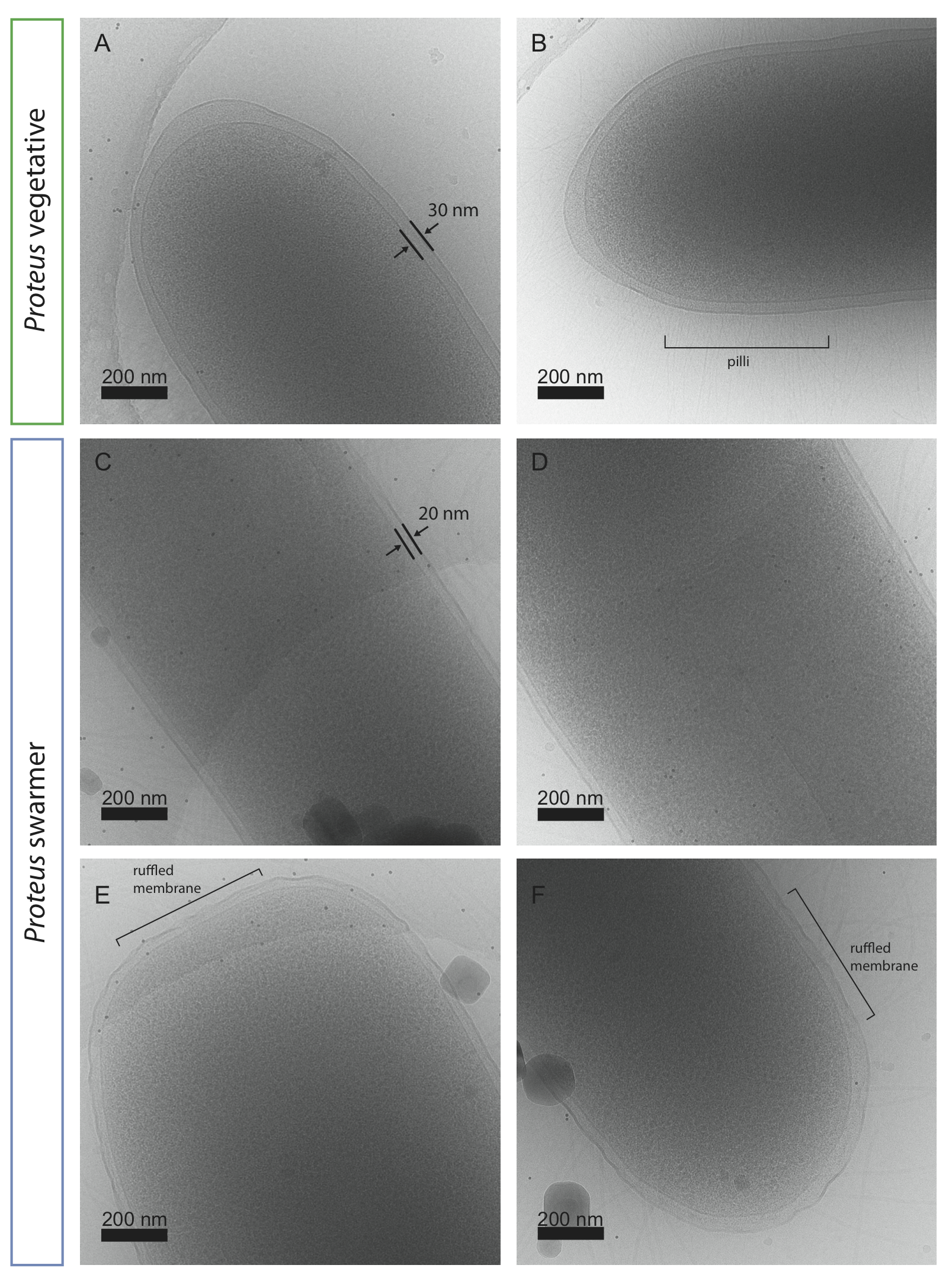

**Fig. S8**

**Figure S9. Electron cryotomography** **of *V. parahaemolyticus* vegetative and swarmer cells reveals cell membrane alterations and defects.** *V. parahaemolyticus* vegetative cells (A, B) have a smooth outer membrane that we also observed in *V. parahaemolyticus* swarmer cells (C); however some swarmer cells exhibited membrane blebs and vesicles (D-F). Ruptures in the cell wall (C) and variable cell diameter (E, F) were also observed. We were unable to determine if these features were the result of increased sensitivity to blotting pressures used when freezing the grids (D-F). Cells are suspended over a thiin carbon film containing circular holes that appear in some of the images (C, E, F).

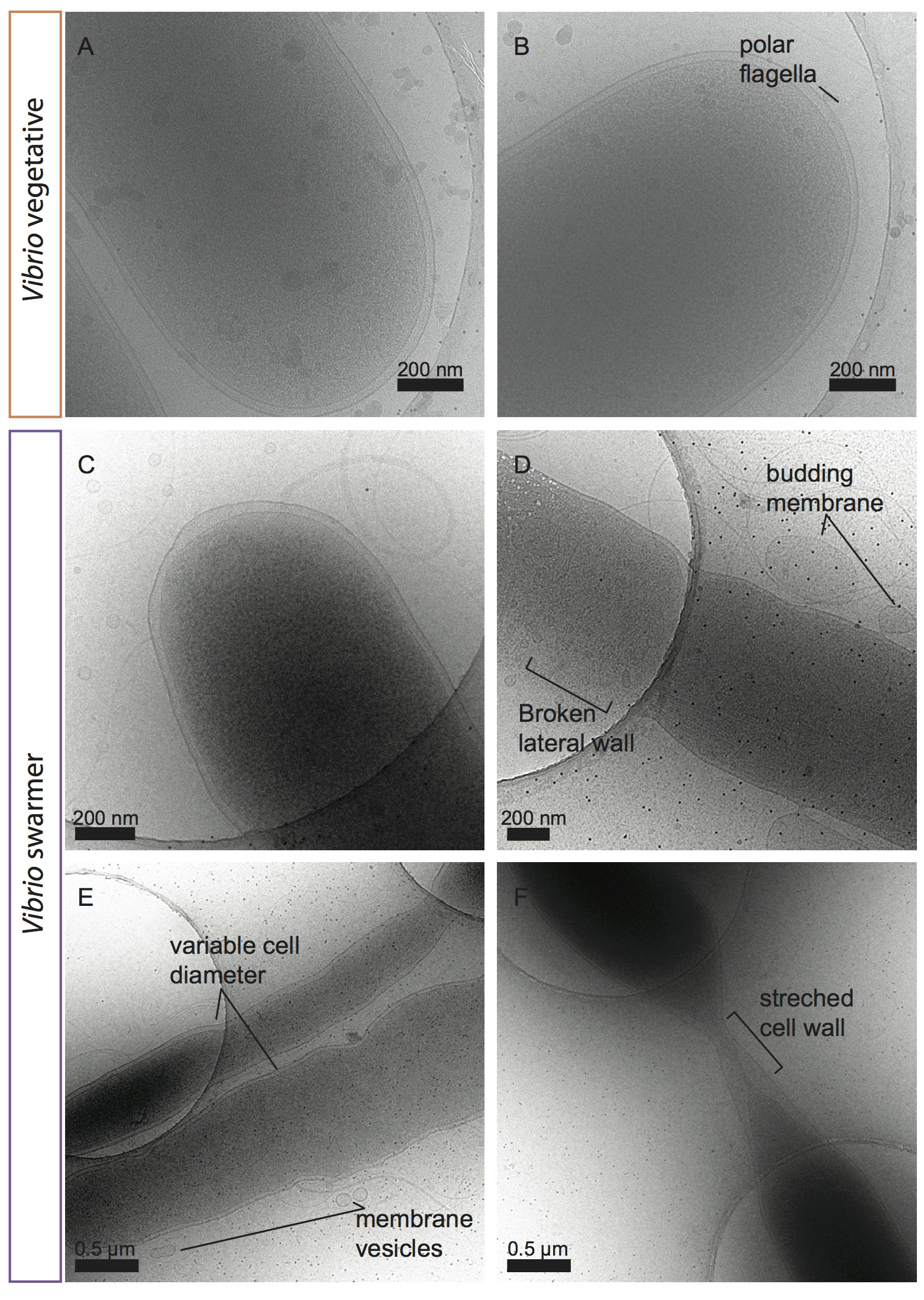

**Fig. S9**

**Figure S10. Sensitivity of cells to treatment with 1x MIC cephalexin.**

A) A phase contrast microscopy image showing *P. mirabilis* and *V. parahaemolyticus* cells treated with 1X MIC cephalexin in the microfluidic device shown in Fig S12. Time = 0 min depicts the time point cells immediately after the introducing cephalexin (1X MIC) to cells. At time = 180 min, we observed a higher frequency of swarmer cells compared to vegetative cells. Dead cells lose their phase contrast (they go from appearing dark to light/transparent). Membrane blebbing and filamentation are a direct result of cephalexin treatment. Scale bar = 10 μm.
B) A cartoon depicting the time course of cells treated with cephalexin. Time progresses from left to right. The ring structure represents the division protein FtsZ.

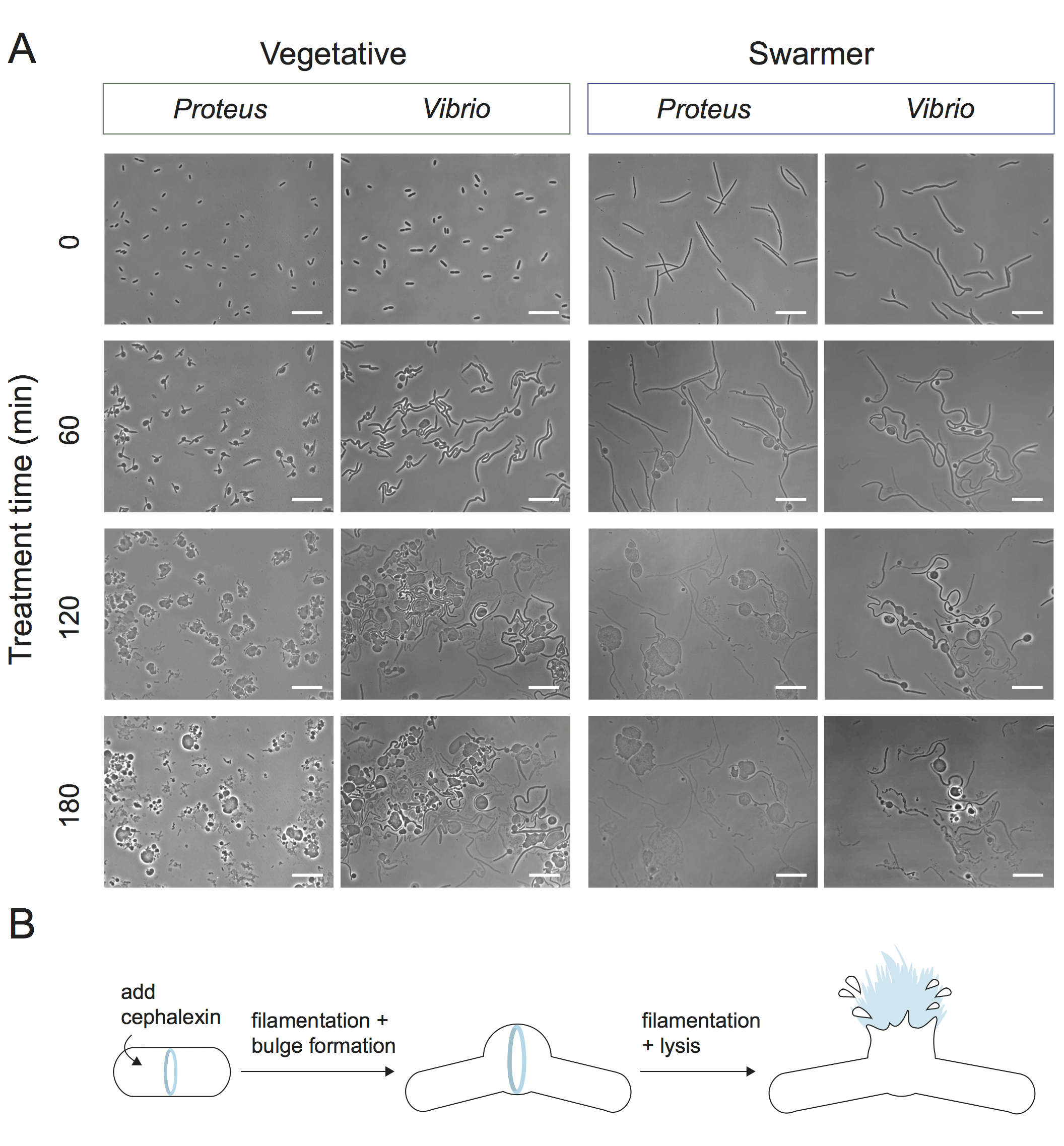

**Fig. S10**
**Figure S11. No significant change in the growth rate of cells treated with β-lactams.** Vegetative and swarmer cells do not show a significant change in growth rate when treated with β-lactam antibiotics. We monitored the growth rate of vegetative exponential phase cells and swarmer cells for 15 min in the microfluidic growth device described in Figure S12. A) A plot of cell growth in the presence of cephalexin (1X MIC); and B) a plot of cell growth in the presence of penicillin G (1X MIC) [*n* > 40 cells, from at least 2 independent experiments]. The box plot depicts the median, 1^st^ and 3^rd^ quartiles (“hinges”), and the 95% confidence interval of the median (“notches”). Black dots represent outliers in data.

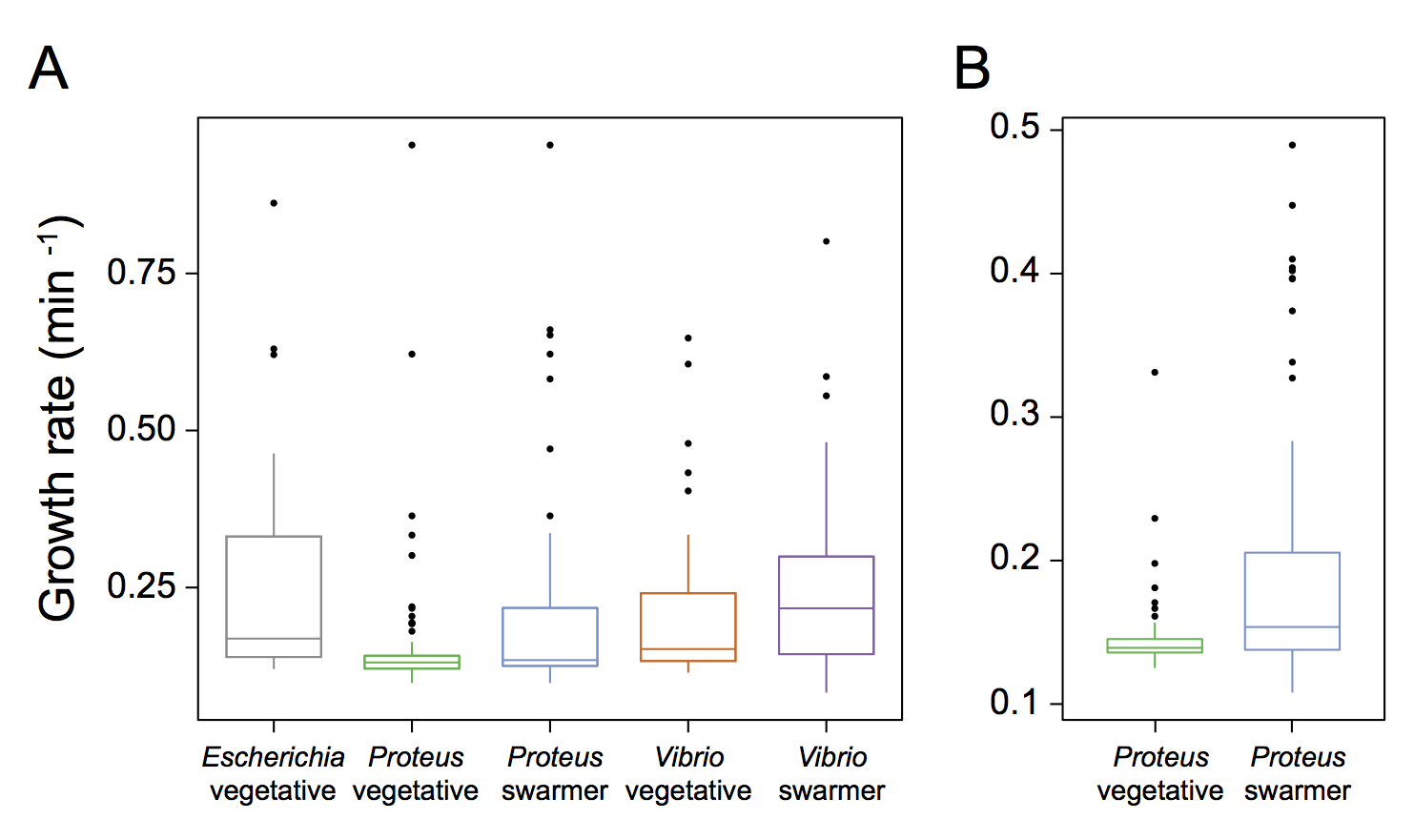

**Fig. S11**

**Figure S12. A cartoon depicting the structure of the microfluidic flow device used in measurements of swarmer cells response to antibiotics.** A 250 μM thick layer of PDMS was applied to a cover glass. A 6 mm x 4 mm rectangular section of PDMS was removed using a scalpel and replaced with a nutrient media containing 2% (w/v) a warm solution of agarose. After the agarose had solidified and cooled, we pipetted 2 μL of a suspension of cells on the agarose surface, waited until the excess liquid was absorbed by the agarose, and inverted the agarose sandwiching the bacteria between the glass coverslip and agarose. The PDMS flow chamber was placed directly onto the PDMS coated coverglass ensuring the agarose pad was centered in the flow channel. A constant flow of nutrient media at 20 μL/min was supplied to the device using a syringe pump. Cell growth was monitored at 30 °C. Images were collected every 1 min for 3 h.

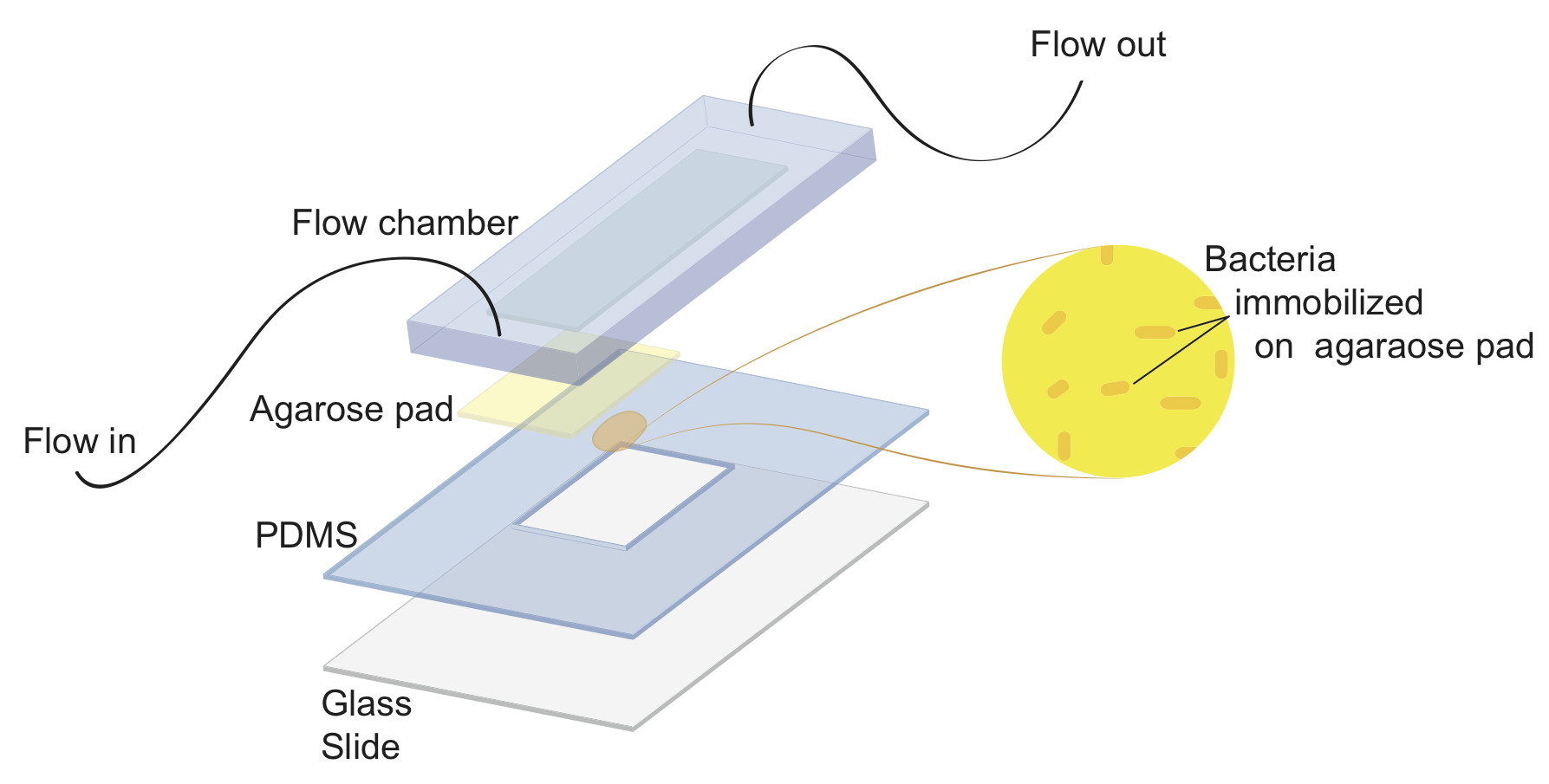

**Fig. S12**

**Figure S13. Gravity flow setup used with the reloadable cell bending microfluidic device.** An image of the gravity flow system we used for precisely delivering fluids at user-defined flow rates for bending measurements. We assembled the flow system next to an inverted microscope; a VC-6 channel valve controller is located outside of image. See methods for operation of the system.

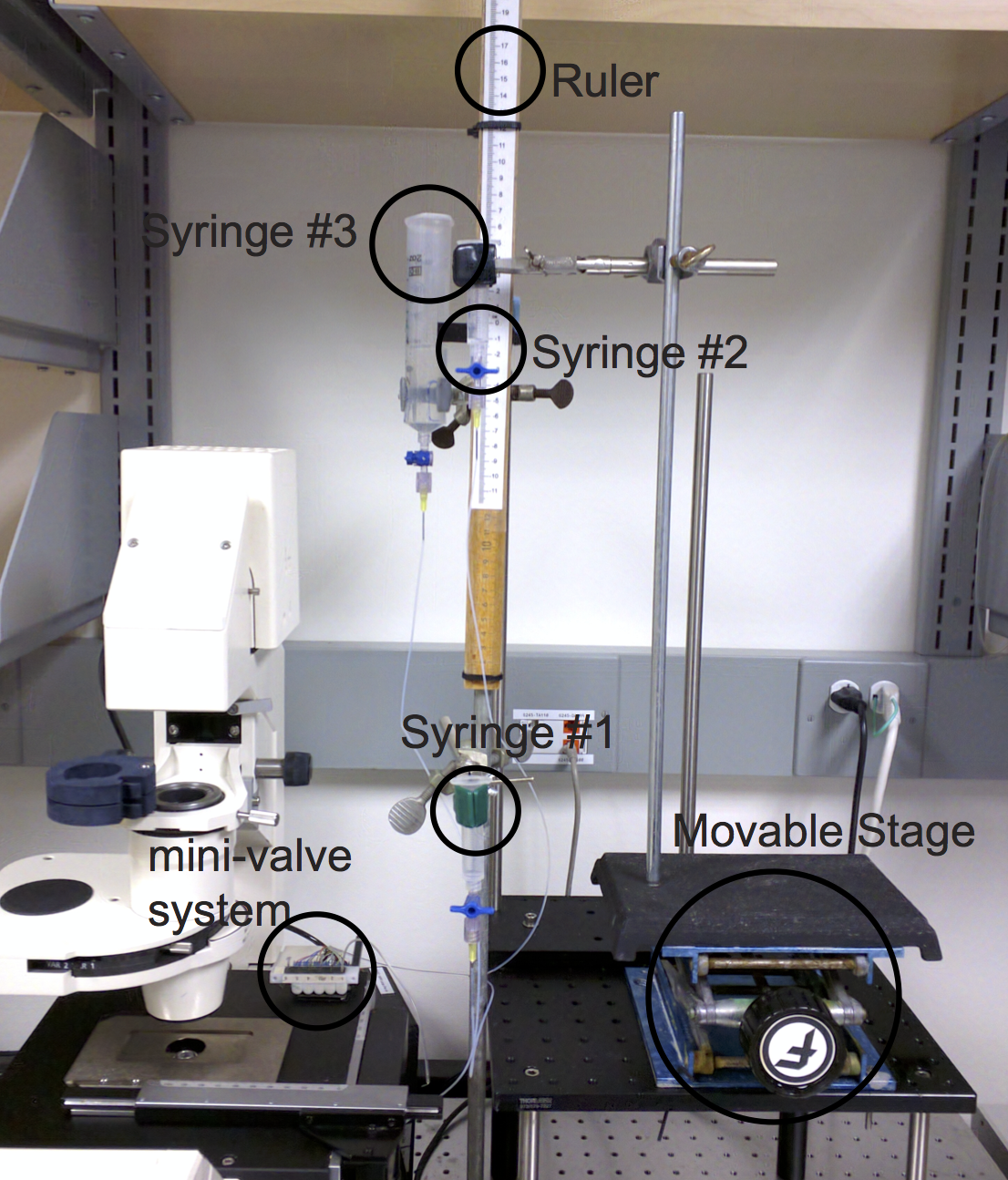

**Fig. S13**

**TABLE**

**Table S1. Muropeptides analyzed by UPLC-MS in positive ion mode.** Identification of muropeptides by MS. We identified muropeptides within (± 0.01 m/z) of the calculated muropeptide mass values. Identified peaks corresponded to mass values with following adducts (1H^+^, 2H^+^, and 3H^+^). MS peaks were used to identify and quantify area under curve of UPLC UV_205nm_ peaks (Figure S8).

| Peak | Retention time (min) | Calculated mass | | | Length of stem peptides |
| --- | --- | --- | --- | --- | --- |
|  |  | H+ | 2+ | 3+ |  |
| 1 | 7.7 | 871.3784 | 436.1892 | 291.1261 | Tri |
| 2 | 8.6 | 928.3999 | 464.6999 | 310.1333 | Tetra-Gly(4) |
| 3 | 9.2 | 699.2936 | 350.1468 | 233.7645 | Di |
| 4 | 10.8 | 999.4370 | 500.2185 | 333.8123 | Penta-Gly (4) |
| 5 | 11.6 | 1243.5202 | 622.2601 | 415.5067 | Tri-Tri (-DS) |
| 6 | 12 | 942.4155 | 471.7078 | 314.8052 | Tetra |
| 7a | 13.6 | 1723.7102 | 862.3551 | 575.2367 | Tri-Tri |
| 7b | 20.6 | 1723.7102 | 862.3551 | 575.2367 | Tri-Tri |
| 8 | 21.3 | 1851.7970 | 926.3985 | 617.9323 | Tetra-Tetra-Gly(4) |
| 9a | 21.7 | 1794.7756 | 897.8878 | 598.9252 | Tetra-Tri |
| 10 | 22.6 | 1922.8341 | 961.9171 | 641.6114 | Tetra-Tetra-Gly (5) |
| 9b | 22.8 | 1794.7756 | 897.8878 | 598.9252 | Tetra-Tri |
| 11 | 24.1 | 1865.8127 | 933.4063 | 622.6042 | Tetra-Tetra |
| 12 | 28.6 | 2846.2313 | 1423.6156 | 949.4104 | Penta-Gly(5)-Tetra-Tetra |
| 13 | 30.1 | 2789.2098 | 1395.1049 | 930.4033 | Tetra-Tetra-Tetra |
| 14a | 30.7 | 1774.7402 | 887.8701 | 592.2467 | anhydro Tetra-Tri |
| 14b | 32.1 | 1774.7402 | 887.8701 | 592.2467 | anhydro Tetra-Tri |
| 15 | 33.2 | 1845.7865 | 923.3932 | 615.9288 | anhydro Tetra-Tetra |

**DERIVATIONS**

**Derivation of the equation(s) describing the behavior of a rod-shaped bacterium bending under laminar fluid flow**

The model for bending a cell is based on the mechanics model of a suspended rod (cantilever) bending under its own weight (Fig. S14). Analytical solutions have been developed for this model, but are not adequate to describe the large deformations measured in our setup.

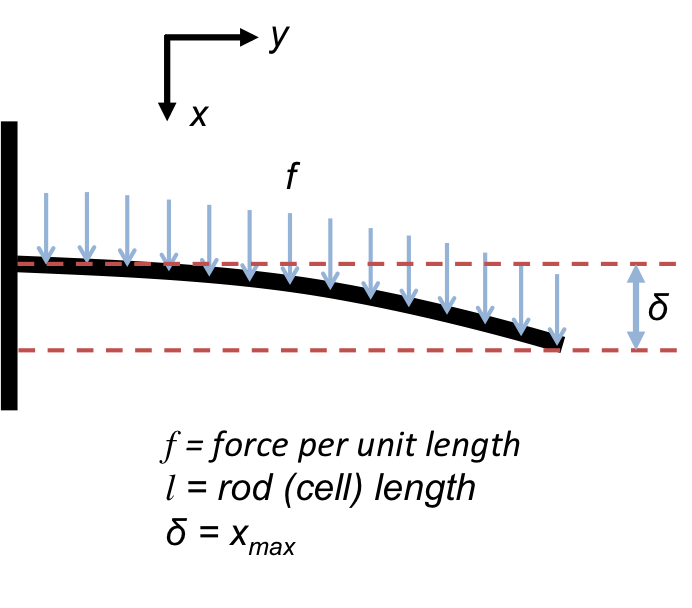

**Figure S14.** Schematic model of a cantilever or cell bending under its own weight.

**General bending of cantilevered rod equation**

At each point along the cantilevered rod or cell, there is an external stress and an internal stress. The external stress is the application of force due to gravity or forces from the fluid. The internal stress, or response, comes from the mechanical properties of the rod, which is what we are interested in probing. By setting these two stresses equal to each other, we can relate the two and extract meaningful information about the material properties of the rod. First we will consider how the internal stress is calculated.

*Internal Stress*:

It is helpful to consider that along every point of the rod there is an associated curvature, *κ*, defined as the inverse of the radius, *R*, formed by a circle at that point. The curvature is related to the infinitesimal angle, Δ*θ*, and infinitesimal arc length, Δ*y*, at the point of interest (see Fig. S15).

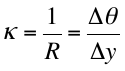
(1)

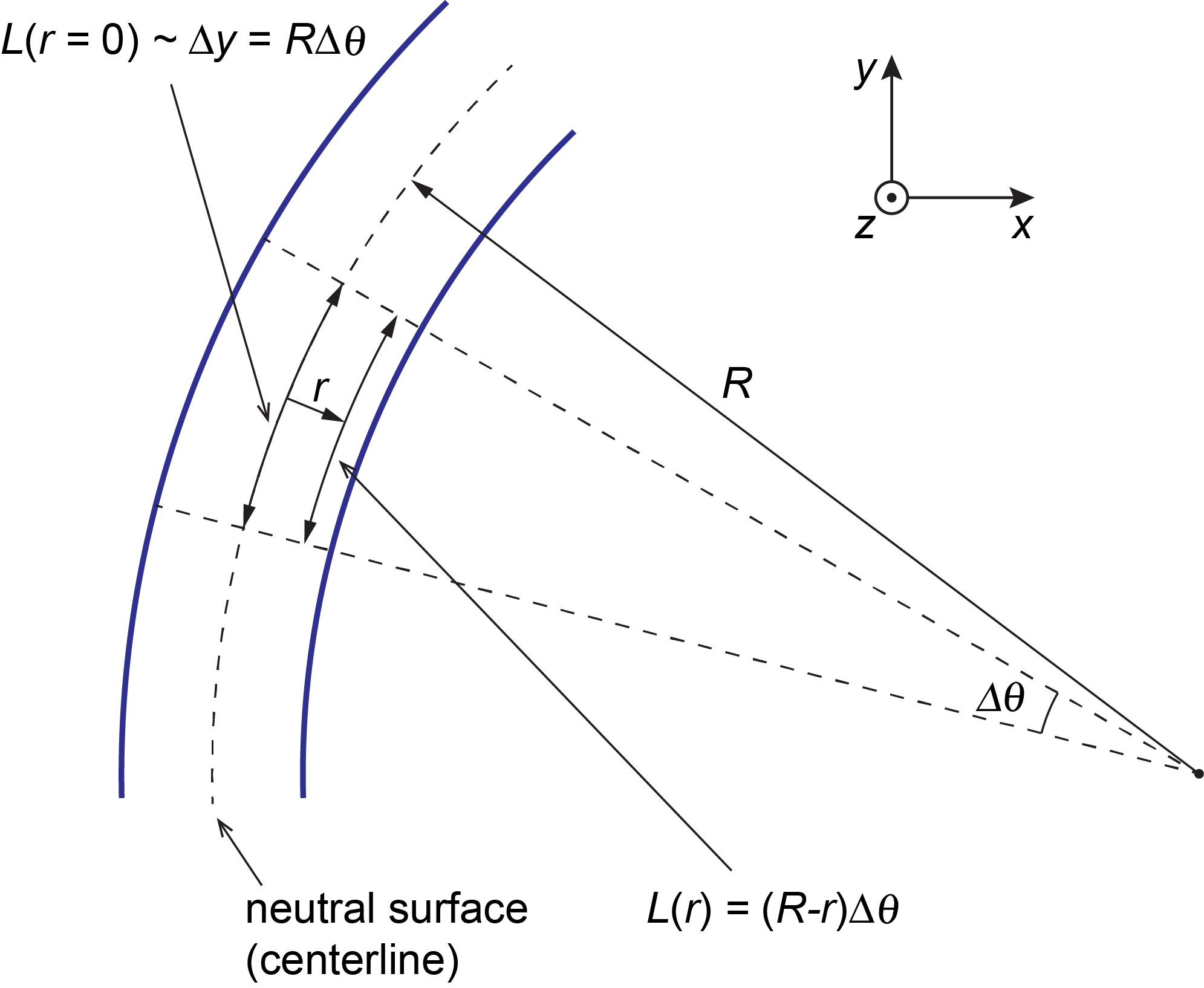

**Figure S15.** Schematic of a curved geometry at each point along the rod (cell). Note that moving away from the centerline by a distance, *r*, shortens the arc length made by Δ*θ*. The image is rotated with respect to Figure S14 in order to reflect the final geometry we are interested in but the coordinate system is the same.

The extensional strain, *ε*, is the ratio of extension and length and is defined as follows at each point within the cross-section:

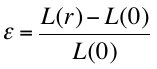
(2)

We can substitute the information from Figure S16 into Equation 2:

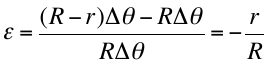
(3)

Equation 3 relates extensional strain to the local radius of curvature, *R*, and the position within the rod (cell) cross-section, *r*. Tensile strain, *σ*, is the force per unit area and is related to extensional strain by the Young’s Modulus, *Y*:

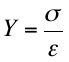
(4)

which traditionally has units of Pascals (Pa), or N m^-2^. Substituting Equation 3 into 4:

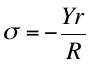
(5)

Equation 5 relates the tensile strain to Young’s Modulus, position in the cross-section, *r*, and local radius of curvature, *R*. Assuming a linearly elastic material, the total internal force in a particular cross section must be equal to zero and is, therefore, the summation of all tensile strains multiplied by the total area:

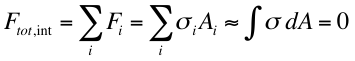
(6)

The total internal bending moment, *M*, however, is not zero. This is because the directional aspect of the bending moment, *r*, counteracts the negative sign of the tensile strain when moving from compression to extension (i.e. in Figure S16 moving from the right part of the rod to the left). The *total* internal bending moment for a cross-section is defined as:

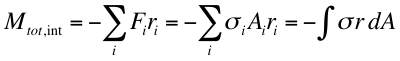
(7)

Substituting in equation 5:

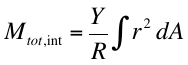
(8)

The area moment of inertia (second moment of inertia), *I*, is defined as:

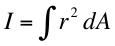
(9)

Substituting Equation 9 into Equation 8 gives us:

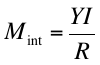
(10)

Equation 10 describes the moment of inertia of a cross-section of the cell. For clarity, we rewrite it as a function of position in the *y*-direction (see Figs. S15 & S16 for coordinate system).

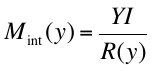
(11)

The area moment of inertia for a hollow disc cross-section can be derived to:

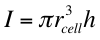
(12)

where *h* is the thickness of the disc (cell wall thickness) and *r*_cell_ is the radius of the rod (cell). This quantity, *I*, will be useful later when calculating the Young’s Modulus of the cell wall from the flexural rigidity (*YI*).

*External Stress*:

Equation 11 provides a relationship between the local internal bending moment, the flexural rigidity, and the radius of curvature within the rod (cell) for a given cross-section. However, these are internal stresses. The external stresses have to be balanced, meaning we have to write the bending moment for the external force on the rod (cell) at each point. For now, we will assume that force is evenly distributed along the rod (cell).

At any point along the cell, *y*, the external bending moment, *M*_ext_, (Equation 15) is the total load of the force along the remainder of the cell (Equation 13) times the location of the centroid of that force,
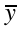
, (Equation 14):

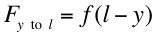
(13)

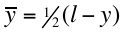
(14)

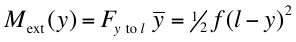
(15)

where *f* is the force per unit length and *l* is the length of the rod (Figure S16). Note that the bending moment is really a torque at that particular point along the rod.

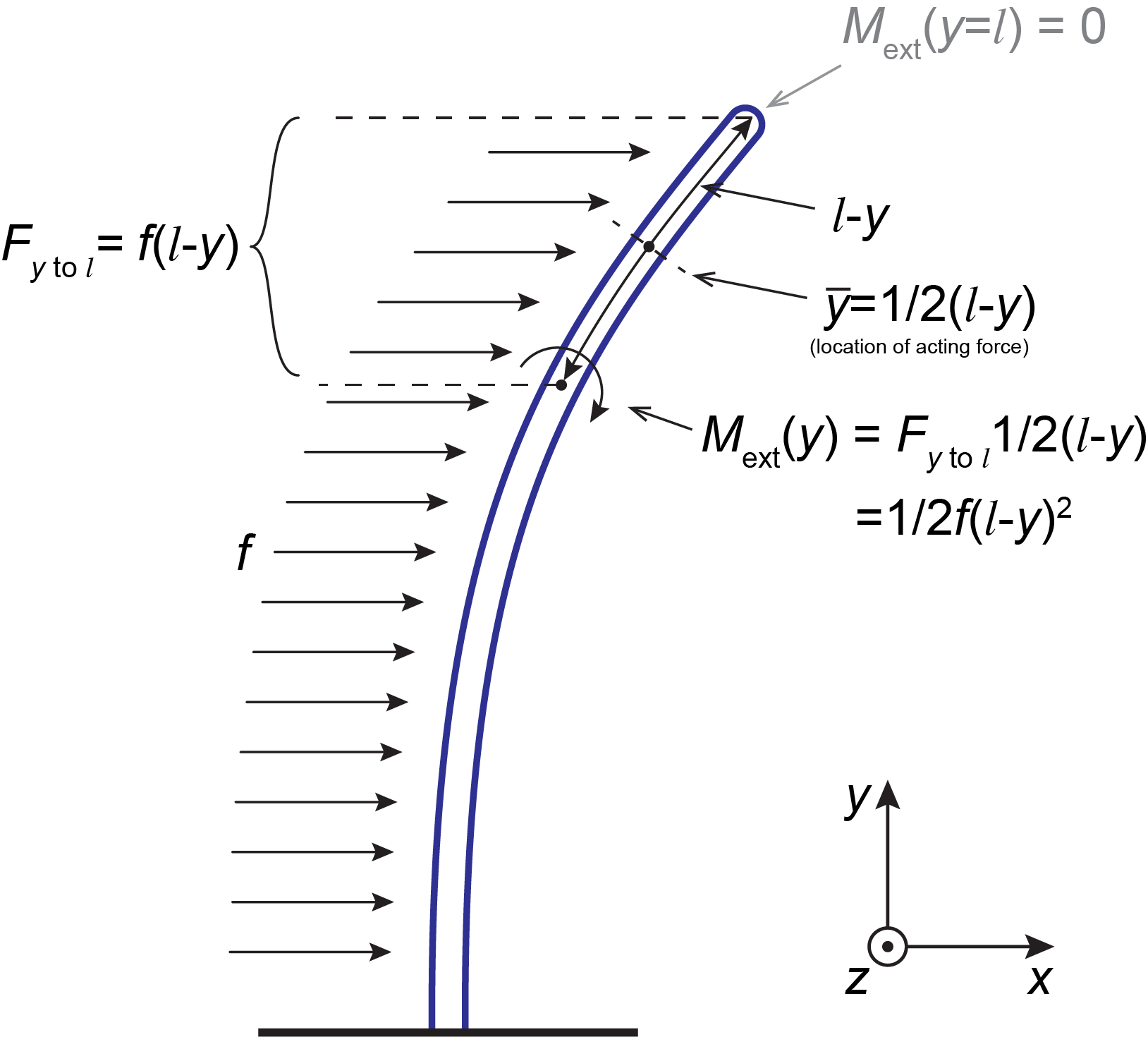

**Figure S16.** Schematic showing the derivation of the external bending moment for a cantilevered rod (cell) with a constant force, *f*, along the rod. The bending moment shown, *M*(*y*), is for any arbitrary point along the cell and goes to zero at the tip (when *y* = *l*).

Now we balance the internal and external bending moments at each point along the rod (*M*_int_(*y*) = *M*_ext_(*y*)) and solve for the curvature, yielding the equation:

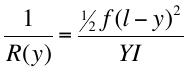
(16)

We are interested in the *x*-position (deflection) of the rod at each point along *y*. This can be accomplished by realizing 1/*R*(*y*) is the local curvature and the curvature of any function is defined as:

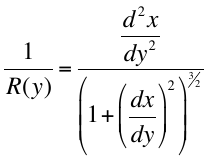
(17)

where here we implement the derivative with respect to *x* (1/*R*(*y*) and 1/*R*(*x*) are equivalent) in order to get a final function with respect to *x*. Setting equation 16 and 17 equal to each other yields the general differential equation:

$\frac{d^{2}y}{{dx}^{2}}={\frac{M_{\mathrm{ext}}}{YI}\left( 1+\left( \frac{dy}{dx} \right)^{2} \right)}^{\frac{3}{2}}$	(18)

where:

$M_{\mathrm{ext}}=\frac{1}{2}f\left( l-y \right)^{2}$	(19)

Assuming the first derivative is zero (appropriate *only* for small deflections), yields the following function of *y*.

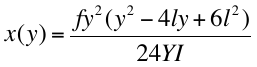
(20)

The maximal deflection, *x*_max_, is at the tip, where *y* = *l*, yielding the equation:

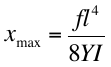
(21)

Equation 21 describes the maximal tip deflection in a rod (cell) with very small deformations and a constant force along the cell. This equation was derived in the supplementary material from Amir et al.^1^

**Derivation of models with fewer assumptions:**

The geometry of our system (Fig. S16) is quite similar to what is shown in Fig. S15, but there are three main factors that must be taken into account when modeling at large deformations:

(1) The solution up to this point (Equations 19, 20, & 21) does not take into account the arc length of the cell as it bends. In other words, the analytical solution makes it seem that when the cell bends significantly, the length of the cell actually gets longer.
(2) In a laminar flow system, the force (*f*) along the cell is not uniform; it changes based on the laminar flow profile.
(3) There must be some sort of attenuation of the force on the cell as it bends substantially and its orientation with respect to the flow becomes non-perpendicular. *This effect is taken into account in the B2 and C2 models and is discussed later.*

(1) *Addressing the arc length:*

We need a model that takes into account the fact that the tip of the cell moves downward (-*y* direction) as the cell is bent substantially. What is needed is a term that takes into account the arc length of the cell, *s*, which, for any differentiable function starting from *y* = 0 is:

$s(y)=\int_{0}^{y} \sqrt{1+\frac{dx}{dy}}dy$	(22)

Replacing *s* with *y* in the bending moment in Equation 19 gets us:

$M_{\mathrm{ext}}(y)=\frac{1}{2}f\left( l-s\left( y \right) \right)^{2}$	(23)

Addition of the arc length into the bending moment means the differential equation (Equation 18) is no longer amenable to an analytical solution. It will need to be solved numerically. Equations 18 & 23 are the basis for the model (B) used to calculate maximum deflections of cells in the microfluidic device. The numerical solution is discussed later in this document.

(2) *Addressing the non-uniform laminar flow profile:*

The laminar flow profile (described by the Poiseuille flow equation) provides the velocity vector along the geometry of the device. Hence, *f* is actually a function of a number of parameters including the characteristic flow velocity, *v^*^*, chamber width, *w*, chamber height, *h*, as well as cross-sectional area of the cell π.

$v=v^{*}\left[ \frac{z}{2h}\left( 1-\frac{z}{h} \right)-4\sum_{n=0}^{\infty} \frac{\sin\left[ \left( 2n+1 \right)\frac{\pi z}{h} \right]}{{(2n+1)}^{3}\pi^{3}}\left( \frac{\cosh\left[ \left( 2n+1 \right)\pi\left( \frac{y}{h}-\frac{w}{2h} \right) \right]}{\cosh\left[ (2n+1)\frac{\pi w}{2h} \right]} \right) \right]$	(24)

$f(y)=4\pi\eta r\left. \frac{\partial v}{\partial z} \right|_{y, z=r}$	(25)

where *η* is the viscosity and *r* is the radius of the cell. The partial derivative of velocity with respect to *z* in Equation 25 is calculated at a given *y*-position and at a *z*-position close to the surface (e.g. where *z* equals the radius of the cell).^2^ We use a *z*-position of 0.5 μm in all of our calculations. Note that a practical value of *n* for the summation in Equation 24 is approximately 10, which is the default value in our calculations.

This complicated flow function makes the model for cell-bending untenable to an analytical solution—although Amir, et al. did use an approximate constant force for their calculations to obtain their results.

It is important to note that the relationships shown in Figure S17 are true only when there are small deformations and the force along the rod is uniform. In order to take into account a non-uniform force, we must calculate the bending moment along the cell at each point. Ignoring, for now, the arc length, the general definition of the bending moment is:

$M_{\mathrm{ext}}\left( y^{'} \right)=F_{y^{'}\mathrm{to} l}y_{c}\left( y^{'} \right)=\int_{y'}^{l} f(y)dy\frac{\int_{y'}^{l} f(y)\left( y-y^{'} \right)dy}{\int_{y'}^{l} f(y)dy}=\int_{y'}^{l} f(y)\left( y-y^{'} \right)dy$	(25)

where *y’* is referring to the point at which the bending moment is being calculated and *y_c_* is the centroid force coordinate (effective position of the applied force).

If *f*(*y*) is assumed to be constant, then Equation 25 simplifies to:

$M_{\mathrm{ext}}\left( y^{'} \right)=f\int_{y'}^{l} \left( y-y^{'} \right)dy=\frac{1}{2}f{(l-y')}^{2}$,	(26)

and is the same result as in Equation 15 (model **A1**), which we got by inspection. In our system, however, the force is not constant along the length of the cell and the force must remain a function of *y*, as in Equation 25. The addition of an arc length further complicates the equation:

$M_{\mathrm{ext}}\left( y^{'} \right)=\int_{s'}^{l} f(y)\left( s-s' \right)ds$	(27)

where *s’* is the arc length at *y’*. Equations 18 & 27 describe the model called **C1** and its numerical solution will be discussed later.

*Addressing the attenuation of force as it becomes parallel to the flow:*

If there are large deformations, the angle the cell makes with the fluid flow will not be perpendicular at all points along the cell. In fact, longer cells readily fold over onto themselves completely. The parts of the cell that are parallel to the flow no longer meet the standard criterion of Stoke’s drag on a rod perpendicular to the flow. Thus, *f* is also a function of the local angle the cell makes with the fluid flow. In the extreme case, we can assume that parts of the cell that are parallel to the flow will exhibit no shear force.

A way to implement this type of attenuation is to multiply *f* by a scaling coefficient between 0 and 1, where 1 is used for parts of the cell that are completely perpendicular to the fluid flow and encounter the full force of the flow, and 0 is used for parts that are completely parallel and encounter no force.

$A(y)=\frac{2}{\pi}{tan}^{-1}\left( \left. \frac{dy}{dx} \right|_{y} \right)$	(28)

which is derived by the cell having a particular angle, *φ*, at every point along the cell with respect to the *y*-axis.

$\frac{dy}{dx}=tan\varphi$	(29)

so that when phi equals π/2 (90°), that part of the cell is parallel to the *y*-axis and when it is zero, it is parallel to the *x*-axis. Dividing by π/2 (arctan(∞) = π/2) gives us a coefficient between 0 and 1 that attenuates the force based on angle of the cell.

We can include the attenuation factor to Equation 27:

$M_{\mathrm{ext}}(y^{'})=\int_{s'}^{l} f(y)A(y)\left( s-s' \right)ds$	(30)

where *s’* is *s* at *y’*. Equations 18 & 30 describe the model called **C2** and its numerical solution will be discussed later.

**Models and Numerical solution methods:**

The various models and governing equations are presented schematically in Figure S17. They are solved using a procedure file written for Igor Pro 6.37 (Wavemetrics, Inc.).

A-models (not used in manuscript):

The ‘A’ models are governed by Equations 20 and 21 and are not full solutions to the differential equation (Equation 18) because the solutions disregard the first derivative and simplify the bending moment. Furthermore, the forces applied are approximate. These models are not used for fitting and are for illustration purposes only.

*Model A1:* A1 is governed by Equation 21 and follows a simple fourth-order exponential.

*Model A2:* A2 is governed by Equation 20, but the arc length is numerically calculated using Equation 22. The *x*_max_ value is found where the cell length, *l*, equals the arc length.

B-models:

The B-models are governed by the full differential in Equation 18, but the forces in the bending moment are not fully integrated as shown in Equation 25. For the ‘B’ models, the force for a bending moment at a particular point along the cell is applied exactly on-half the distance (arc length) to the end of the cell at *y_c_*. The bending moment is:

$M_{\mathrm{ext}}\left( y \right)=\frac{1}{2}f(y_{c}){(l-s(y))}^{2}$,	(31)

where

$y_{c}=y @ \frac{1}{2}\left( l-s\left( y \right) \right)+s(y)$.	(32)

The value *y_c_* is calculated numerically. When force attenuation is taken into account, Equation 31 becomes:

$M_{\mathrm{ext}}\left( y \right)=\frac{1}{2}f(y_{c})A(y_{c}){(l-s(y))}^{2}$,	(33)

which is a simplification of Equation 30. A more accurate measure of the bending moment is addressed in the ‘C’ models.

*Model B1:* B1 is governed by Equations 18, 31, and 32 as explained above. No force attenuation is calculated.

*Model B2:* B2 is governed by Equations 18 and 33. This solution takes into account force attenuation as the cell bends.

C-models:

The ‘C’ models are governed by the full differential equation (Equation 18) as well as the proper definition of the bending moment (Equation 27). The ‘C’ models are preferable to the ‘B’ models due to their strict adherence to the proper bending moment, but the ‘C’ models come at a significantly higher computational price. Much of the higher cost of calculation comes from the need to iterate the bending moment much like the arc length discussed earlier.

The model solves for the cell profile at a given cell length using an iterative process, i.e., a starting cell profile is given, the differential equation is solved, a new starting cell profile given, and so forth, until the differential equation is minimized for a given cell length. Sometimes the “just-calculated” solution (cell profile) can “overcompensate” for the previous one and cause the solutions to oscillate indefinitely between two meta-states without ever converging. Hence, the impact of the newer solution must be attenuated by a factor we call the *solution contribution factor (SCF)*. For C1 and C2, this sometimes needs to be set as low as 2.5%, meaning the latest solution will contribute only 2.5% to values of the previous one. Hence, when using a low SCF value, it takes more iterations to reach the converged cell profile. Changing the SCF value only prevents non-convergence and does not alter the final solution.

*Model C1:* Governed by Equations 18 and 27. No force attenuation is calculated.

*Model C2:* Governed by Equations 18 and 30. This solution takes into account force attenuation as the cell bends.

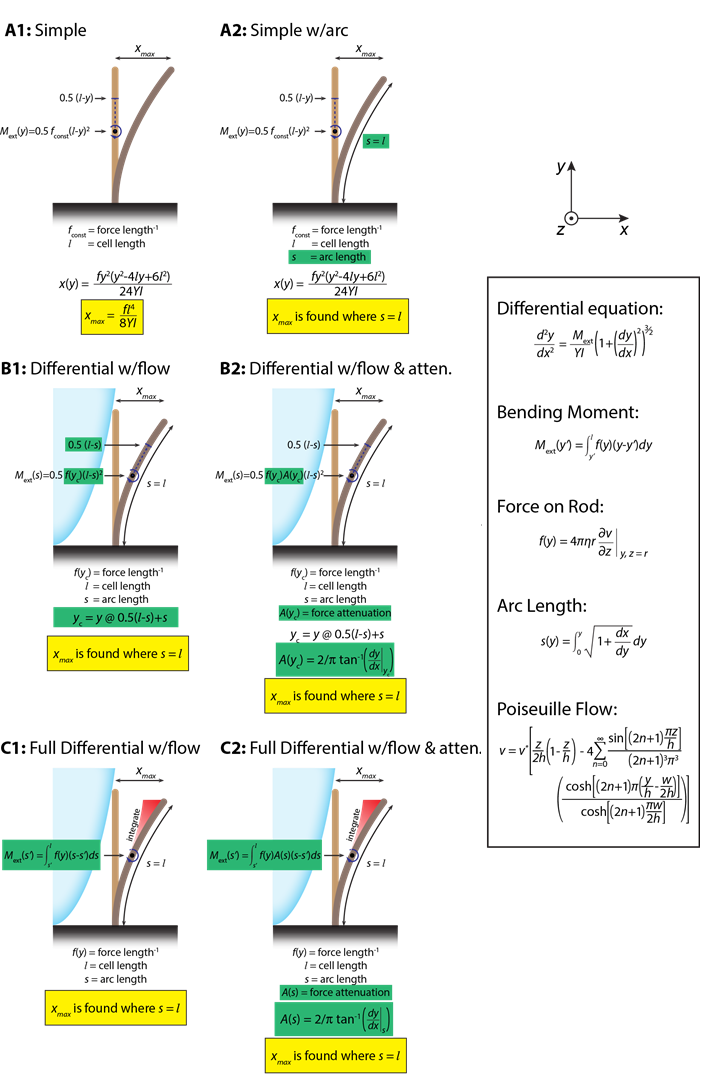

**Figure S17.** Schematic of the models described in this section.

**Comparison of Models**

The resulting displacement versus cell length curves were compared for models B1, B2, C1, and C2 for high (4x10^-20^ N m^2^), medium (1x10^-21^ N m^2^), and low (1x10^-22^ N m^2^) bending rigidities given the experimental conditions in the flow cell and constants as follows:

- *h =* 28.9x10^-6^ m (chamber height)
- *w* = 100x10^-6^ m (chamber width)
- *v** = 0.084252 m/s (characteristic flow velocity)
- *r* = 0.5x10^-6^ m (cell radius)

Figure S18 shows that, qualitatively, the behavior of all four models is very similar. There is a more substantial deviation between the models at higher cell lengths for B2 and C2 due to the effect of the attenuation factor.

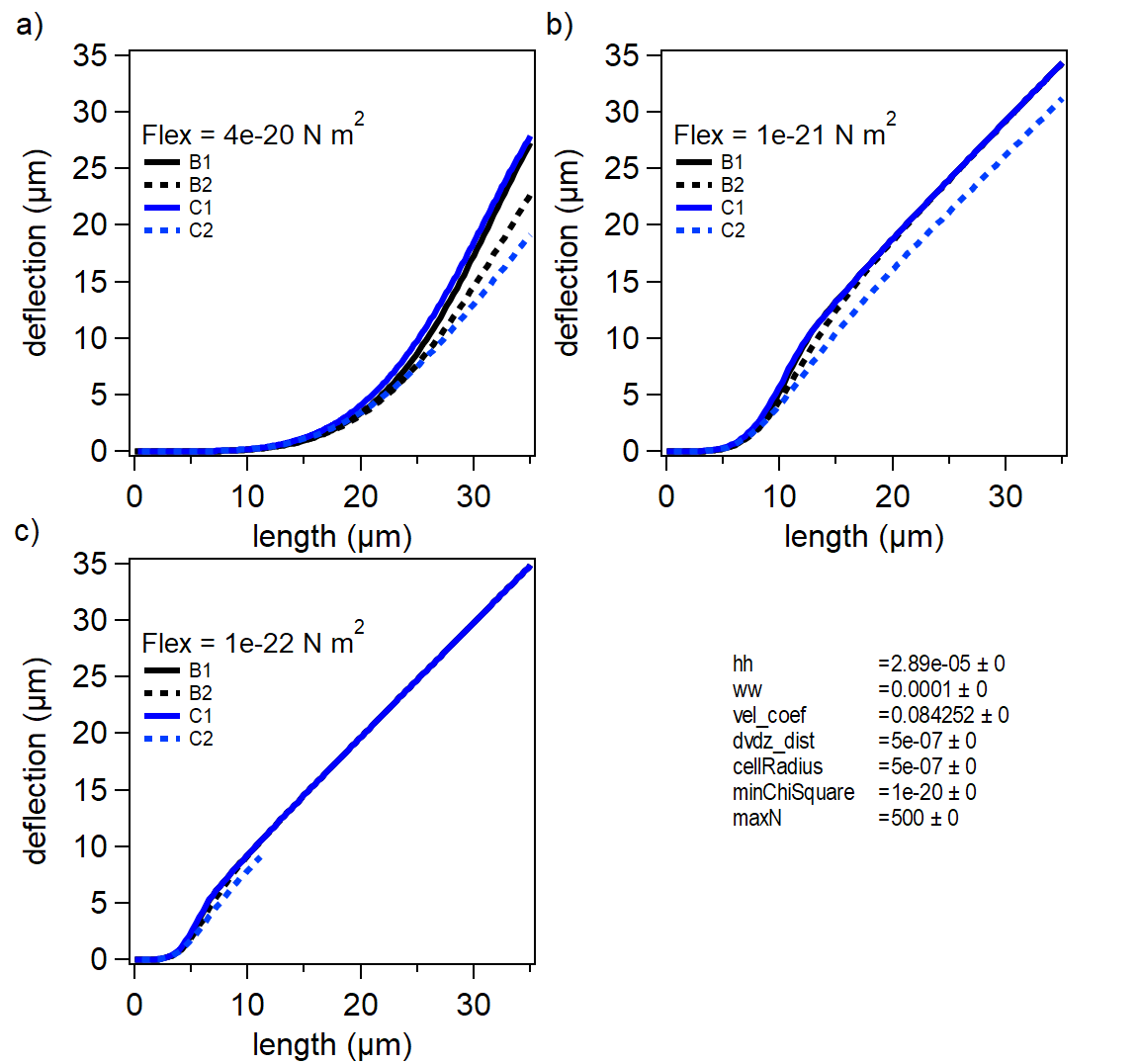

**Figure S18.** Comparison of deflection versus cell length for models B1, B2, C1, and C2 at bending rigidities of a) 4x10^-20^ N m^2^, b) 1x10^-21^ N m^2^, and c) 1x10^-22^ N m^2^.

All datasets were analyzed by models B1, B2, C1, and C2 using a fitting function written into the Igor procedure file (based on a nonlinear least squares regression analysis). As expected, the resulting bending rigidity values vary depending on the model chosen. Model C1, which performs a full integration of the bending moment, estimates the highest bending rigidities, while C2, which is the same as C1 except the force is attenuated as the cell bends, estimates the lowest. Figure S19 shows the results from the four models for each strain. All of the models exhibit the same trends and all converge similarly with low error in the bending rigidity, **but we chose to use an average of C1 and C2 for all reported values**. The C models most closely match the real physics that are applied to the cell. We are averaging the two C models (with and without force attenuation) because we believe that the force attenuation of the C2 model may be an oversimplification of the drop in force as the cell bends with respect to the flow. It is important to note the bending rigidity values calculated for *E. coli* match closely with previous reports.^1,3^

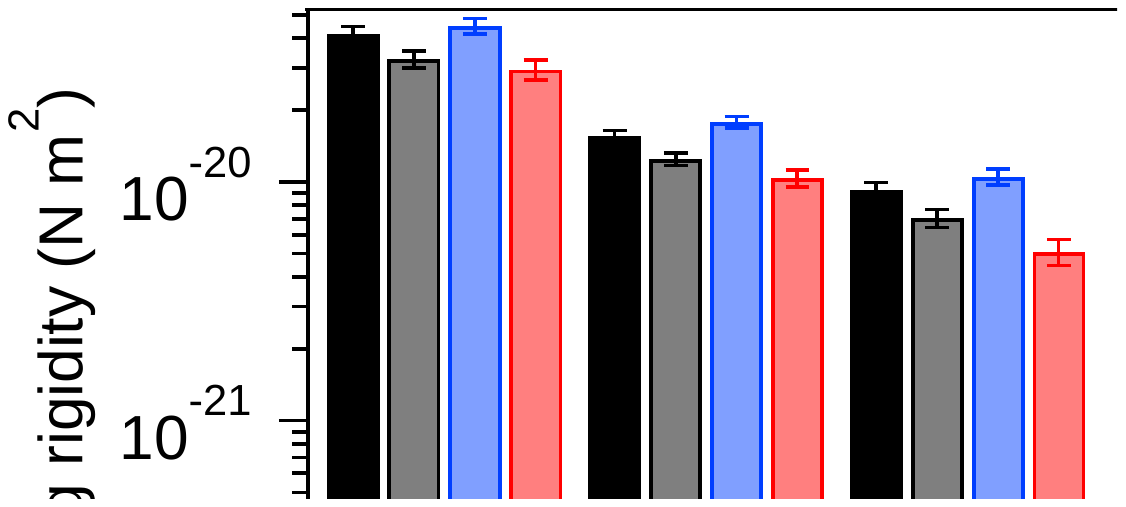

**Figure S19.** Calculated bending rigidities for each strain tested. Error bars indicate 95% confidence intervals from the curve fitting algorithm.

**Effect of Force Attenuation**

To allow the reader to better visualize the effect of force attenuation factor, A(y) (used in the B2 and C2 models), we include plots of the resulting cell profiles in Figure S20. These cell profiles are calculated from the equations described above in the model—the *x*_max_ (displacement length) is calculated from the data that comprise these plots. Here we compare the B1 and B2 models at cell lengths of 10 and 20 µm at the given parameters. For the case where no force attenuation is implemented (B1), the cell has a higher total force placed on it throughout the entire length of the cell. Therefore, the cell “folds over” more readily, meaning the displacement length (*x*_max_) would get closer to the cell length value than for the case where force attenuation is applied used. These trends hold for the C1 and C2 models as well.

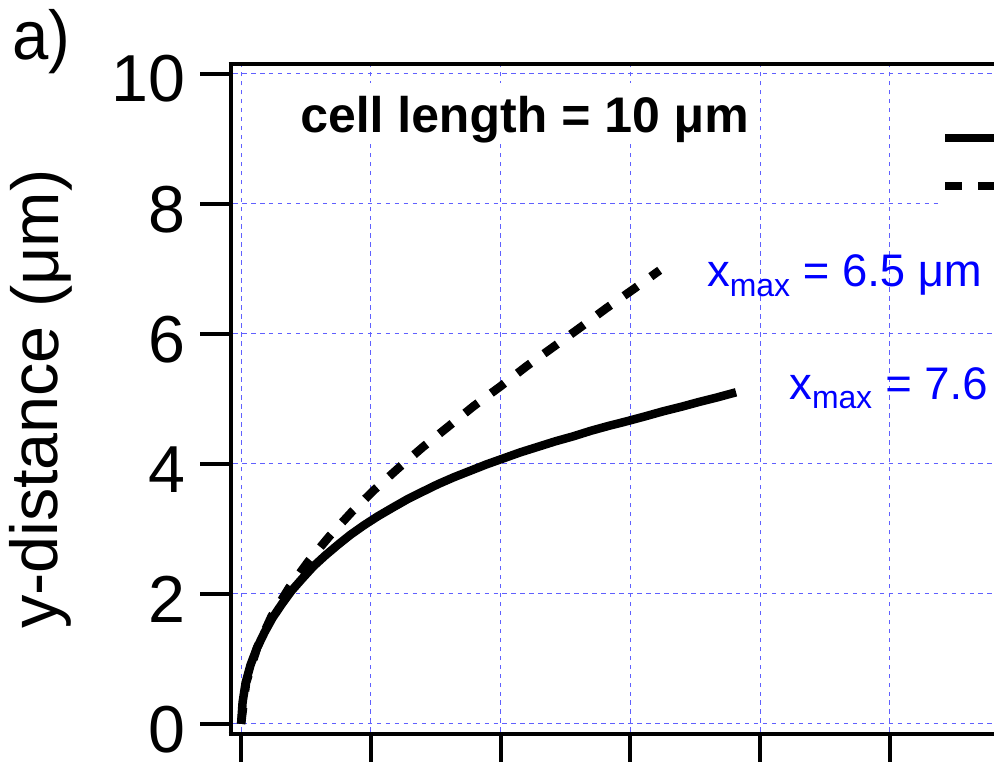

**Figure S20.** Comparison of B1 (without attenuation) and B2 (with attenuation) cell profiles for a) 10 µm and b) 20 µm long cells with the given parameters. Flow is indicated by the red arrow. The B1 model, which does not take into account the attenuation of force that the cell makes as it orients itself perpendicular to the flow, “folds over” more readily, increasing the x_max_ value compared to the B2 model, which take into account the attenuation.

**Effect of Limiting the Data Range for the Curve Fitting Algorithm**

We tested the effect of limiting the horizontal axis data range (i.e., cell length) on the curve fitting algorithm. We fitted the B1 model to the *Proteus* swarmer (with *pflhDC*) dataset and varied the range of fit from the full range (to ~56 µm) down to 5 µm. The fitting results are shown in Figure S21a (zoomed in on Figure S21b) as a function of the fitting range. The vertical dashed lines correspond to the upper limit of the cell length range for each respective fit. The resulting flexural rigidity parameters are shown in Figure S21c. It is clear from Figure S21a & b that the 5 and 10 µm cell length ranges are poor approximations of the full experimental behavior. The data indicates that, if the fit is limited to cell lengths before the linear displacement region dominates (i.e., when the slope becomes ‘1’), the resulting values of bending rigidity can change. Due to the variation in the bending rigidity parameter when limiting the fit to low cell lengths, and the consistent values achieved when extending the fits to longer cell lengths, we decided to fit as much of the cell length as possible for each variant. These trends hold for the other fitting models used in this manuscript.

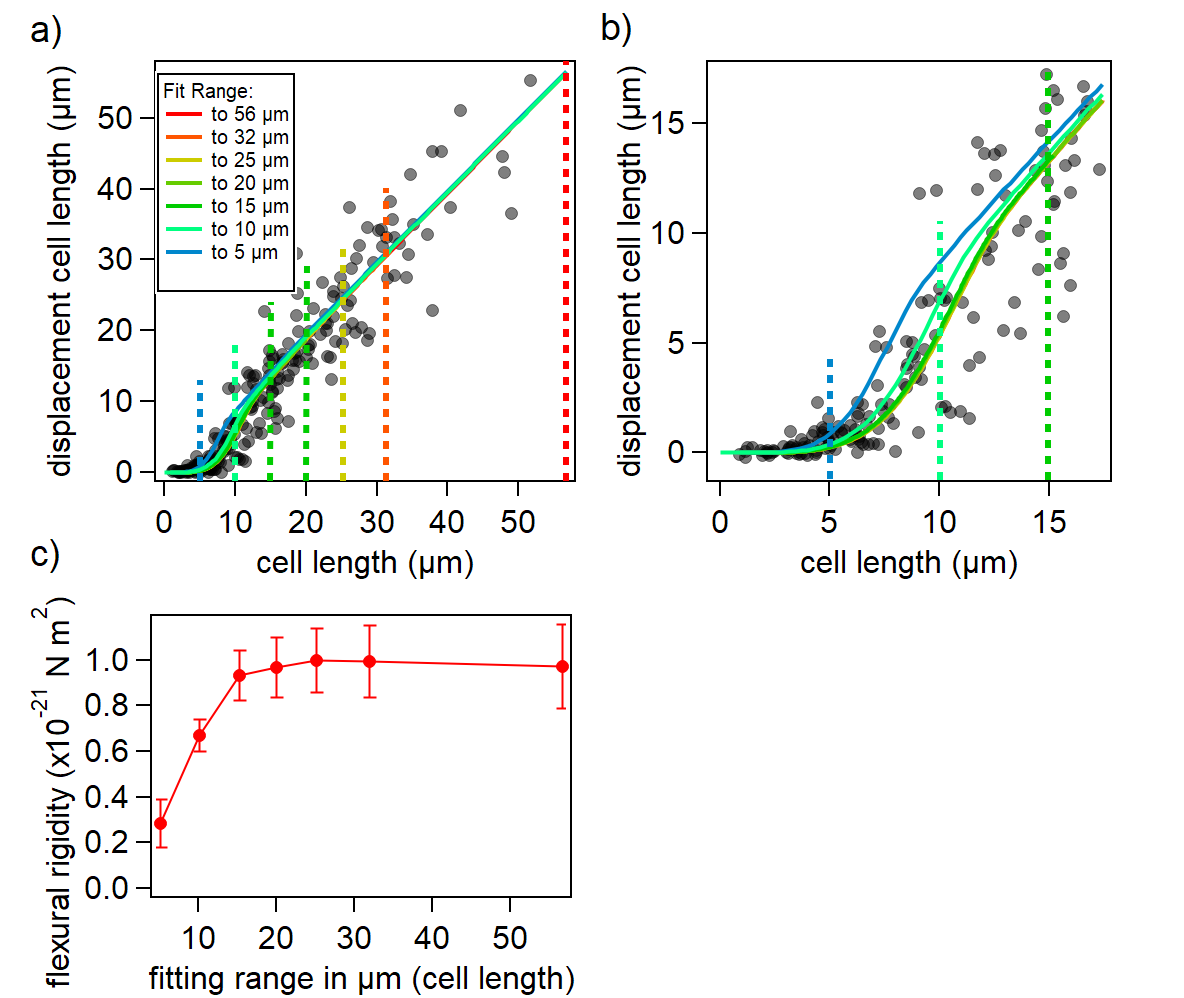

**Figure S21.** Effect of limiting the data range (cell length) on the calculated bending rigidity from the curve fitting algorithm. A) *Proteus* swarmer (plus *pflhDC*) data with fits to the B1 model in ranges from 5 µm to the entire range (~56 µm). The curve fits are extended beyond the fitting range for illustrative purposes. B) Zoomed in plot from (A). C) Bending rigidity versus fitting range in µm. Error bars indicate 95% confidence intervals. These trends hold for the other fitting models used in this manuscript.

**Numerical Considerations for Solving the Model**

To solve the differential equation, the coordinate space and orientation of the cell must be defined. At first, it would seem prudent to solve the differential equation so that the slope at the origin (base of cell) is zero, as in Figure S15. This strategy works quite well until the cell becomes very long or very flexible. This is a problem because, at large deformations, the slope near the end of the cell could become -∞. Alternatively, rotation of the cell counter-clockwise by 90 degrees creates a problem at the origin (i.e. a slope of +∞). This issue is easily solved by rotating the solution between the two extremes at 45 degrees, giving the initial condition of dy/dx = +1 at the origin and a minimum slope of -1 along the cell. The rotation of the coordinate system does not in any way alter the differential equation (Equation 18), since the curvature is indifferent to orientation. However, the solution must be rotated back into normal space as follows:

$y=\frac{x_{rot}}{\sqrt{2}}+\frac{y_{rot}}{\sqrt{2}}$	(33)

$x=\frac{x_{rot}}{\sqrt{2}}-\frac{y_{rot}}{\sqrt{2}}$	(34)

The numerical solution also requires the arc length as a function of *y*. This function, *s*(*y*), will change depending on the solution to the differential equation (hence this is an integro-differential equation). Therefore, an iterative process of solving the differential equation then calculating the arc length, putting it back into the differential equation, and so forth is required until a solution has converged. The convergence is determined by minimization of chi-squared, which is defined as:

$\chi^{2}=\sum\frac{{(current-prevous)}^{2}}{previous}$	(35)

where a typical desired chi-square value is 1x10^-20^ for our system. After convergence, the *x_max_* value is calculated by determining the deflection where the cell length, *l*, equals the arc length.
